# Transcranial Alternating Current Stimulation (tACS) Entrains Alpha Oscillations by Preferential Phase Synchronization of Fast-Spiking Cortical Neurons to Stimulation Waveform

**DOI:** 10.1101/563163

**Authors:** Ehsan Negahbani, Iain M. Stitt, Marshall Davey, Thien T. Doan, Moritz Dannhauer, Anna C. Hoover, Angel V. Peterchev, Susanne Radtke-Schuller, Flavio Fröhlich

## Abstract

Modeling studies predict that transcranial alternating current stimulation (tACS) entrains brain oscillations, yet direct examination has been lacking or potentially contaminated by stimulation artefact. Here we first demonstrate how the posterior parietal cortex drives primary visual cortex and thalamic LP in the alpha-band in head-fixed awake ferrets. The spike-field synchrony is maximum within alpha frequency, and more prominent for narrow-spiking neurons than broad-spiking ones. Guided by a validated model of electric field distribution, we produced electric fields comparable to those in humans and primates (< 0.5 mV/mm). We found evidence to support the model-driven predictions of how tACS entrains neural oscillations as explained by the triangular Arnold tongue pattern. In agreement with the stronger spike-field coupling of narrow-spiking cells, tACS more strongly entrained this cell population. Our findings provide the first *in vivo* evidence of how tACS with electric field amplitudes used in human studies entrains neuronal oscillators.

## Introduction

Transcranial electric stimulation (tES) is a noninvasive brain stimulation modality that delivers weak electric currents of typically up to 2 mA peak amplitude to the scalp (Fröhlich, 2014). Transcranial direct current stimulation (tDCS) and transcranial alternating current stimulation (tACS) are the two most common types of tES, where a constant or sinusoidal current waveform is used for stimulation, respectively. The majority of the stimulation current is shunted by the scalp (Fröhlich, 2016) but a relatively small fraction of the current enters the brain and produces electric fields in the range of 0.2-0.5 mV/mm (Datta et al., 2009; Fröhlich, 2016; Miranda et al., 2006; Sadleir et al., 2010). Recently, controversy has engulfed the field due to the heterogeneity of behavioral findings (Brignani et al., 2013; Liu et al., 2018; Sahlem et al., 2015) and the claim that weak perturbations are not strong enough to entrain networks (Huang et al., 2017; Lafon et al., 2017; Liu et al., 2018; Vöröslakos et al., 2018).

Targeted modulation of cortical oscillations and associated long-lasting cognitive and behavioral functions by tACS have been demonstrated in a number of human studies (Antal et al., 2008; Boyle and Frohlich, 2013; Helfrich et al., 2014; Herrmann et al., 2016a; Kasten and Herrmann, 2017; Neuling et al., 2013; Polanía et al., 2012; Vossen et al., 2015; Zaehle et al., 2010). In addition we have recently reported several clinical trials of tACS for the treatment of schizophrenia (Ahn et al., 2019), chronic pain (Ahn et al., 2018) and major depressive disorder (Alexander et al., 2018). At the level of large scale neuronal populations, animal studies have provided evidence for modulation of neural oscillators by tACS *in vitro* and *in vivo* (Ali et al., 2013; Deans et al., 2007a; Fröhlich and McCormick, 2010; Ozen et al., 2010; Radman et al., 2007; Reato et al., 2010a; Schmidt et al., 2014). Finally at the single neuron level, the tACS polarizes the neurons with an alternating change in the direction of polarity resulting in a subthreshold resonance where the hyperpolarization-activated cation current plays a key role in neural response (Aspart et al., 2018; Toloza et al., 2017). Despite these successful demonstrations of weak periodic electric fields interacting with neural oscillators, the mechanism of action remains unclear and debated. Using a computational model of a large-scale cortical network of spiking neurons, we studied (Ali et al., 2013) how tACS entrains ongoing oscillations. We found triangular regions of high-synchrony between stimulation and endogenous neuronal oscillations in a two-dimensional space composed of stimulation frequency and amplitude. The triangular regions are known as Arnold tongues, and were centered on the frequency of endogenous oscillations. The Arnold tongues arise when studying the dynamical properties of coupled oscillators and characterize the regions in parameter space where phase locking appears between two oscillators (Pikovsky et al., 2003).These model-driven predictions along with the existing entrainment hypothesis that is supported by *in vitro* or *in vivo* studies in anesthetized rodents comprise today’s mechanistic understanding of tACS effects. Despite demonstration of long-lasting tACS effects in human studies, the full mapping of the stimulation parameters space and demonstration of resulting synchronization in the form of Arnold tongue structures remains unexplored. The major roadblock to explore this gap in human studies is associated with technical challenges in simultaneous tACS and EEG recording, despite some recent attempts to avoid or filter the large electrical artefact of tACS (Grossman et al., 2017; Negahbani et al., 2018; Witkowski et al., 2016).

In this study, we investigated large-scale neural entrainment of alpha oscillations by tACS in awake head-fixed ferrets. We used tACS amplitudes that induced computationally estimated electric field amplitude of <0.5 mV/mm, which are comparable to the ones for the typical tACS paradigms in humans. We first demonstrate the presence of alpha oscillations (11-17 Hz) in the awake head-fixed ferret in simultaneous recordings from posterior parietal cortex (PPC), primary visual cortex (VC) and LP/Pulvinar complex of thalamus (LP), which point to a top-down control of the network by alpha oscillations, in functional agreement with human studies of alpha oscillations. The alpha amplitude was strongest in PPC which influenced both VC and LP regions in alpha frequency band. Further analysis on putatively classified neurons indicated that the alpha synchrony between field oscillations and firing of narrow-spiking neurons were stronger when compared to broad-spiking neurons. To probe for the presence of an Arnold tongue, the hypothesized mechanism of action of tACS, we systematically varied both stimulation amplitude and frequency. We found evidence supportive of the Arnold tongue for PPC neurons, with narrow-spiking neurons showing stronger phase locking to tACS when compared to broad-spiking neurons. Despite the comparable amplitude of tACS electric field in all three studies region, we did not find Arnold tongue regions for VC and LP neurons. We finally examined the effect of tACS in a computational model of the thalamo-cortical network, which confirmed the main features of our experimental findings. Our work provides in-vivo evidences on how weak electric fields (< 0.5 mV/mm) can entrain neuronal activity, and supports the model-driven predictions about how tACS entrains ongoing neuronal oscillations as demonstrated by Arnold tongue pattern.

## Results

### Endogenous alpha-band oscillations in awake head-fixed ferrets

Understanding the endogenous network dynamics is a necessary step to identify the stimulation targets before examining their engagement by tACS, since we hypothesize that the effect of tACS is a result of the synergistic interaction of endogenous activity and exogenous electric field. We first characterized the endogenous oscillations to validate the existence of previously reported alpha oscillations in the thalamocortical system of awake head-fixed ferrets (Stitt et al., 2018). LFP signals were acquired by low-pass filtering (<300 Hz) the extracellular broadband recordings (Fig. 2 a). Spectral analysis of cranial screw EEG and LFP signals in PPC, VC, and LP showed an alpha peak (12-16 Hz) in PPC, VC, and LP and EEG recordings for all three animals (Fig. 2 b). The alpha oscillations were strongest in PPC and VC, with slight differences in individual alpha peak frequencies between animals: 14.42, 12.32, 14.42 Hz for animals 1, 2, and 3 respectively. Prominent theta oscillations (3-5 Hz) were evident in LP for all animals. To analyze the endogenous oscillations at neuronal level, we extracted the extracellular spikes (n=1539, 701, 391 units in PPC, VC and LP respectively), and clustered the unit waveforms based on spike length (*spkL*) defined as trough-to-peak time. We identified two populations of neurons in PPC (narrow spiking, *n*=689: *spkL=0.361±0.005* ms, broad spiking, *n*=850: *spkL=0.810±0.005* ms) and VC regions (narrow spiking, *n*=416: *spkL=0.283±0.005* ms, broad spiking, *n*=285: *spkL=0.651±0.009* ms), and only one population in LP region (n=391, *spkL=0.642±0.008*) (Fig. 2. c, d). We next examined how different neurons from each region interacted with each other or with the meso- and macroscale neuronal signals (LFP and EEG, respectively).

**Figure 1.**
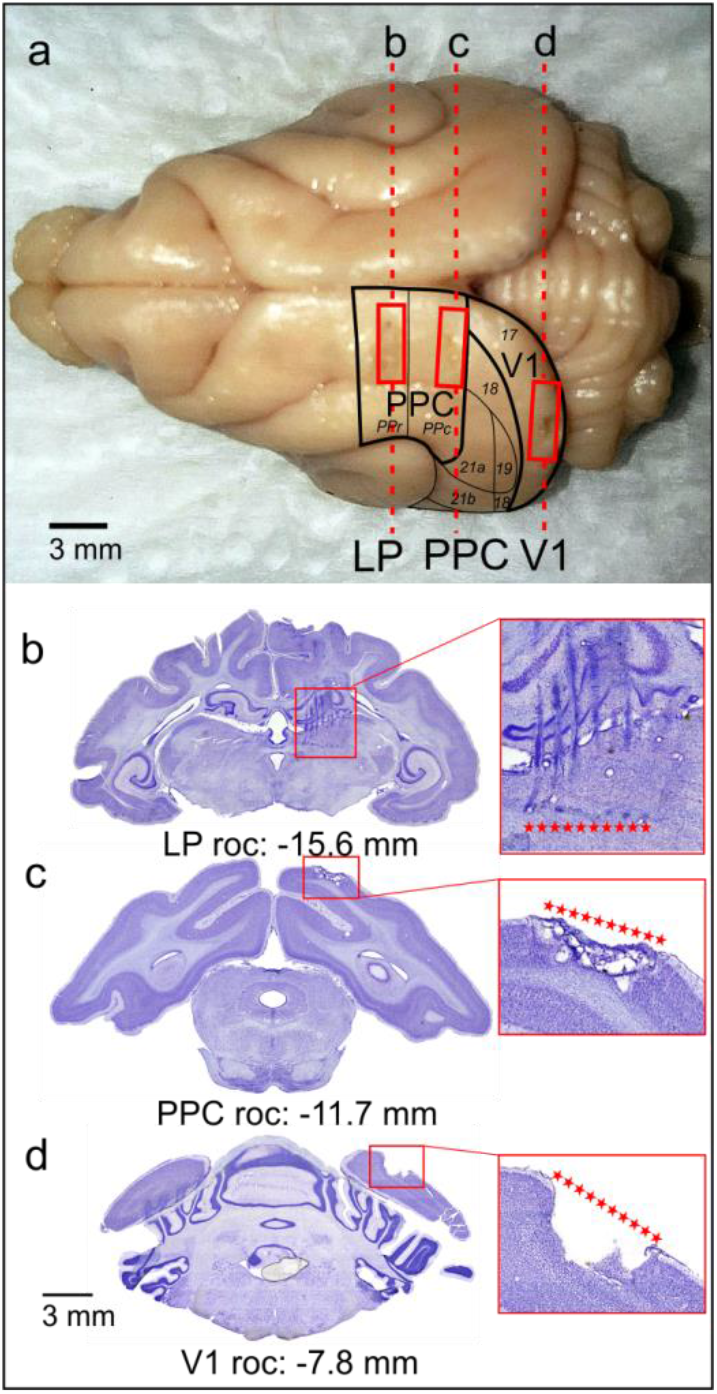
Histological verification of implant locations. (a) Ferret brain showing marks (marked by rectangles) left by microelectrode array implants in LP, PPC, and VC regions. Outlines of cortical fields are based on the ferret atlas (Radtke-Schuller, 2018). Dashed lines indicate the location of cell-stained coronal sections through the center of the multi-electrode array implanted sites shown below (b-d). Corresponding anterior-posterior atlas coordinates are indicated below each section in mm, relative to occipital crest (roc). (b) Section through LP region shows implant electrode tracks and endpoints (dark spots of cell agglomerations above red stars). (c) Section through PPC implant region demonstrates the damage of tissue at the site of implant (below red starts). (d) Section shows the location of VC implant marked by tissue loss (below red starts) caused by the removal of the implant during brain extraction.

**Figure 2.**
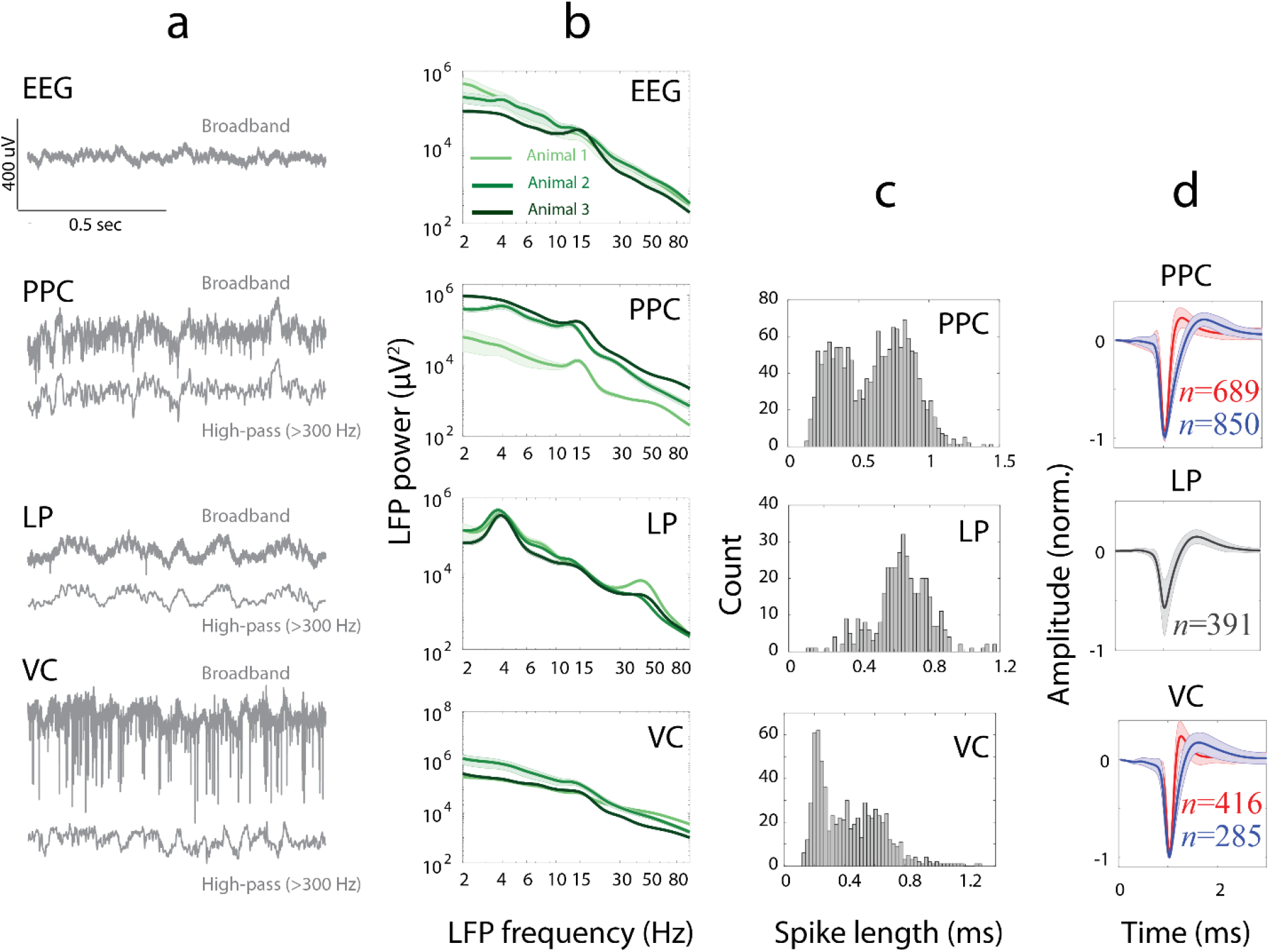
Endogenous alpha-band oscillations in awake head-fixed ferrets. (**a**) Sample 1-sec length EEG, broad-band extracellular recordings and local field potential signals (LFPs) from posterior parietal cortex (PPC), visual cortex (VC) and lateral posterior pulvinar (LP) from a head-fixed awake ferret without performing any task. (**b**) Spectral analysis of EEG and LFPs from three animals, showing the spectrum (mean±SEM) in shades of green with error-bars in light background. The scalp EEG signal demonstrates weak alpha band activity for all animals. PPC shows alpha band activity (12-16 Hz) for all animals. LP shows prominent theta, gamma (35-50 Hz) and a weak alpha activity for all animals. A weak alpha-band activity is evident on primary visual cortex (VC) for all three animals. A closer look at PPC spectra indicates slight difference at individual alpha frequency between animals (14.42, 12.32, 14.42 Hz for animals 1, 2, and 3 respectively). (**c**) Histogram of spike length defined as the time from trough to peak in PPC, LP, and VC. (**d**) Color coded spike waveforms (mean±SEM) for two identified clusters of narrow-spiking (red, *n*=689, 416 in PPC and VC respectively) and broad-spiking (blue, *n*=850, 285 in PPC and VC respectively) neurons in PPC and VC. LP neurons were comprised of only one cluster (n=391).

### Population coupling is stronger for narrow-spiking neurons, and oscillates at alpha frequency band

We recorded broad-band extracellular activity from cortical and thalamic regions and the EEG signal during endogenous oscillations. This recoding enabled us to examine the interaction between neural activities at single-unit and large-scale LFP or EEG levels to better understand the underlying mechanisms of endogenous oscillations. We first asked if there is a relationship between single unit activity and the local dynamics in each region by measuring the population coupling (Okun et al., 2015). We constructed the population firing rate function for each region of interest (ROI) based on its spiking activity. The percentage increase in spike-triggered average population firing rate was extracted at spiking instance compared to the baseline value at 500 ms before spike generation for all narrow- and broad-spiking units in PPC and VC, and for all units in LP. This value provided a measure of how strongly each unit is coupled to local oscillations (Fig. 3 a-c, top row), and related each neurons participation in population code to underlying circuit connectivity (Okun et al., 2015). We observed that the coupling of single units to population rate in all regions oscillates in alpha frequency range, confirmed by spectral analysis. We found prominent peak in the alpha [11-17 Hz] frequency band for all three regions (Fig. 3 ac, bottom row). A weaker theta-band [3-8 Hz] oscillation was also evident in all regions. Comparing the narrow- and broad-spiking neurons, we found that the former had stronger population coupling (Fig. 3 a-c, top row, amplitudes at time lag zero), and the fluctuation of the coupling had more oscillatory power over a wide frequency range including alpha band in both PPC and VC areas (Fig. 3 a-c, bottom row). These results show a population coupling exists for each neuron in recorded cortical and thalamic regions. The population coupling for both narrow- and broad-spiking neurons had an oscillatory pattern with maximum power at alpha frequency band. The population coupling of narrow-spiking neurons oscillated with alpha frequency more strongly than the corresponding measure for broad-spiking neurons.

**Figure 3.**
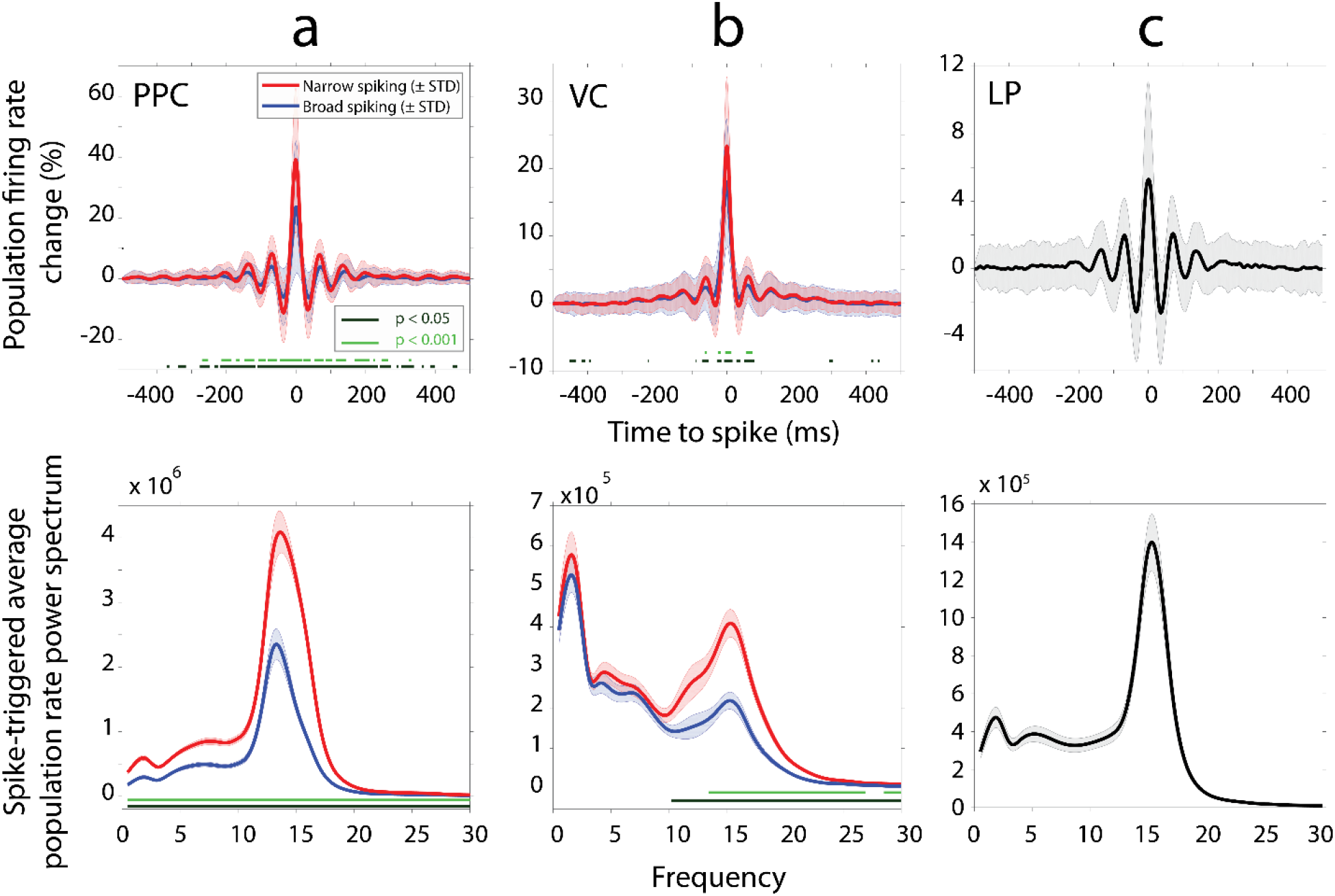
Population coupling is stronger for narrow-spiking neurons, and oscillates at alpha frequency band. **(a, b, c, top)** Percentage change of spike triggered population firing rate (mean ± STD) as a function of time-to-spike. The percentage change is extracted according to corresponding baseline value at t=-500 ms in PPC, VC and LP. The spike-triggered population firing rate functions have larger amplitude at spike instant (time lag zero) for narrow-spiking units (red) compared to broad-spiking cells (blue) in both cortical regions (two-sample t-test at each time point at two significant levels indicated by horizontal green lines). **(a, b, c, bottom)** The power spectrum (mean ± SEM) of spike-triggered population firing rate in PPC, VC and LP. A prominent peak at alpha [11-17 Hz] frequency band is evident for all three regions. The alpha power is higher for narrow-spiking units (red) when compared to broad-spiking units (blue) in cortical regions (two-sample t-test at each frequency point at two significant levels indicated by horizontal green lines).

### Alpha synchronization between EEG and spikes, and prominent engagement of narrow-spiking neurons

Next we examined the alpha synchrony on a larger scale by examining the synchronization between single units in all three ROIs and EEG oscillations. The average spike-triggered EEG showed an oscillatory pattern at alpha frequency band (Fig. 4 a-c, top row). Further analysis showed a non-uniform phase histogram for each region (Fig. 4 a-c, middle row). Narrow-spiking PPC units preferred to spike at 174.67°, confidence interval (CI)=[172.71°, 176.63°] phase of EEG alpha oscillations. The preferred phase for broad-spiking neurons was 173.14°, CI = [170.95°, 175.32°]. For VC neurons the preferred phase was slightly different between narrow (94.06°, CI=[89.50°, 98.62°]) and broad spiking (109.63°, CI=[105.48°, 113.77°]) neurons. The preferred synchronization phase for LP units was 142.63°, CI=[138.44°, 146.82°]. We next extracted the phase locking value (PLV) to examine the strength of spike-to-EEG phase locking as a function of frequency. While the PPC units were phase locked to EEG at a wide frequency range of ~3-32 Hz (Fig. 4 a, bottom), the synchronization was limited to a narrower band of ~10-32 Hz for VC (Fig. 4 b, bottom) and LP units (Fig. 4 c, bottom). Synchronization in the alpha frequency band (12-17 Hz) was evident for single units of all three ROIs. We found the phase-locking between EEG and single units was stronger for narrow-spiking neurons when compared with broad-spiking ones. This behavior was evident in both PPC and VC (Fig. 4 b, c, and bottom). In summary these results show how synchronization is evident between single units and EEG in the alpha (12-17 Hz) band, despite differences in preferred phase between recorded areas. The results also indicate that narrow-spiking units are more engaged in alpha synchronization. We further analyzed the relationship between population coupling of single neurons and their engagement with globally measured alpha oscillations.

**Figure 4.**
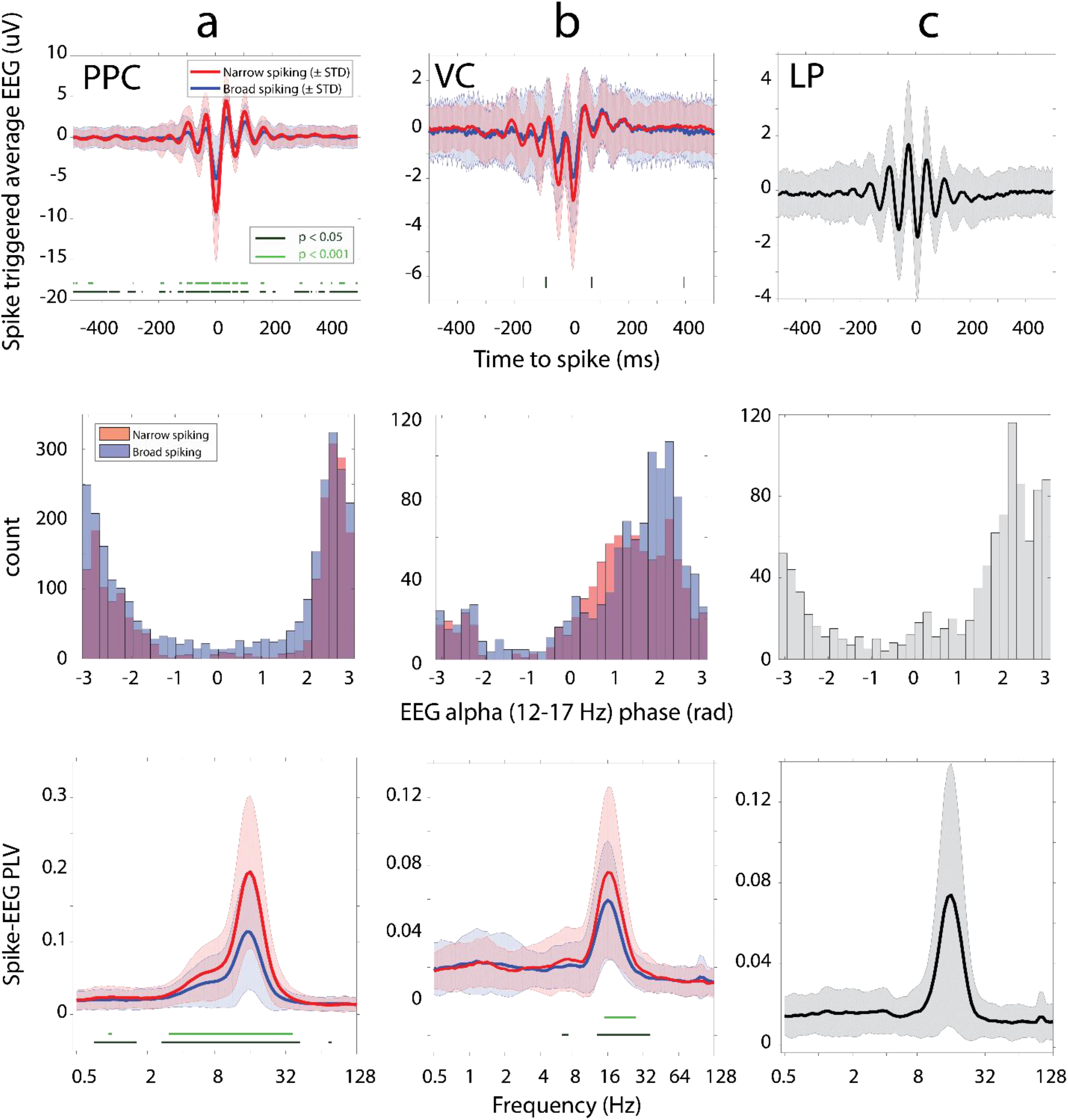
Alpha synchronization between EEG and spikes, and prominent engagement of narrow-spiking neurons. **(a-c, top)** Spike-triggered EEG (mean ± STD) as a function of time-to-spike in PPC, VC, and LP. Spike-triggered average EEG shows an oscillatory pattern for all three regions and shows larger amplitudes at peaks and troughs for narrow-spiking (red) when compared to broad-spiking (blue) neurons in cortical sites (two-sample t-test at each time point at two significant levels indicated by horizontal green lines). **(a, b, c middle)** The phase histogram of spiking units in PPC, VC and LP/Pulvinar related to alpha-band (12-17 Hz) EEG oscillations. Color denotes the counts for narrow- and broad-spiking neurons. Two classes of neurons show similar phase preference in PPC (Broad-spiking neurons: 3.02±0.02 rad, Narrow-spiking neurons: 3.05±0.02 rad), and slightly different phase preference in VC (Broad-spiking neurons: 1.91±0.03 rad, Narrow-spiking neurons: 1.64±0.04 rad). (**a, b, c** bottom) Phase locking between single units and EEG measured by PLV (mean ± SEM) as a function of EEG frequency. Note the synchronization in ~ 3-32 Hz range with prominent peak at alpha (12-17 Hz) frequency range for recorded cortical and thalamic sites. The narrow-spiking cells (red) show significantly higher PLV compared to broad-spiking units (blue) (two-sample t-test at each frequency point with p-values <0.05 indicated by horizontal line).

We found strong correlation between population coupling and engagement by global alpha oscillations in PPC, no correlation in VC and weak correlation in LP (Fig. 5 a-c). In summary these results indicate that the more a PPC unit coupled to population dynamics (chorister), more engaged by globally measured alpha oscillations. Conversely, the less a PPC unit is coupled to local dynamics (soloist), less engaged by globally measured alpha oscillations.

**Figure 5.**
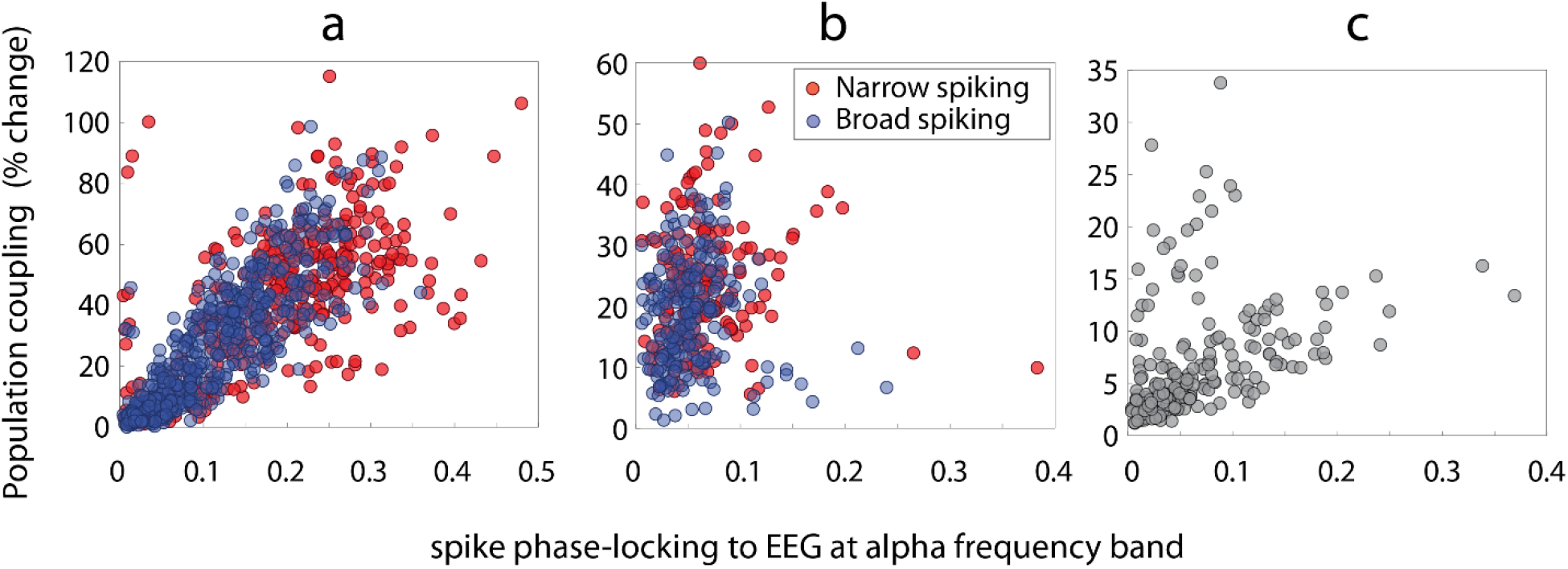
Population coupling of single neurons and their engagement with globally measured alpha correlate strongly in PPC but not in VC and LP. Population coupling of single units versus their engagement by alpha-band EEG is plotted for PPC (a), VC (b), and LP (c). The values on vertical axis are measured by strength of percentage-change in spike-triggered population firing rate at time of spike generation. Color denotes the neuron types. (a) There is a strong correlation between population coupling and large-scale alpha synchronization for narrow-spiking (red, n= 403, r = 0.70, p<0.001) and broad-spiking (blue, n = 582, r = 0.86, p<0.001) in PPC. (b) Population correlation and large-scale alpha synchrony do not correlate in VC (narrow-spiking: red, n=173, r=0.08, p=0.31, and broad-spiking: blue, n=221, r=0.027, p=0.70). (c) Population coupling and long-range synchrony are weakly correlated in LP (n=251, r=0.37, p<0.001).

### Synchronization directed functional connectivity in thalamo–cortical network

To further analyze the dynamics of endogenous network oscillations, we examined the synchronization between all three region pairs using spiking and LFP activities. We first employed the spiking activity to look at the synchronization between recording regions. The population average spike cross-correlations demonstrated synchronous oscillatory structure occurring within alpha frequency band between three region pairs (Fig 6 a-c). The mean power spectrum (±SEM) of cross-correlations indicated a prominent peak in the alpha band (centered on ~15.23 Hz, Fig 6 d-f). These results indicate how spiking activity between PPC, VC, and LP recording sites are synchronized in alpha frequency. We next examined the phase-locking between spikes and the LFP for each region pair to further analyze the between-region synchronization. The spikes from three ROIs were alpha phase-locked to LFP from PPC (Fig. 6 g), VC (Fig. 6 h) and LP (Fig. 6 i). We next computed population average phase locking between LFPs obtained from three regions to understand their interactions (see supplementary figure S1 for individual animals).

**Figure 6.**
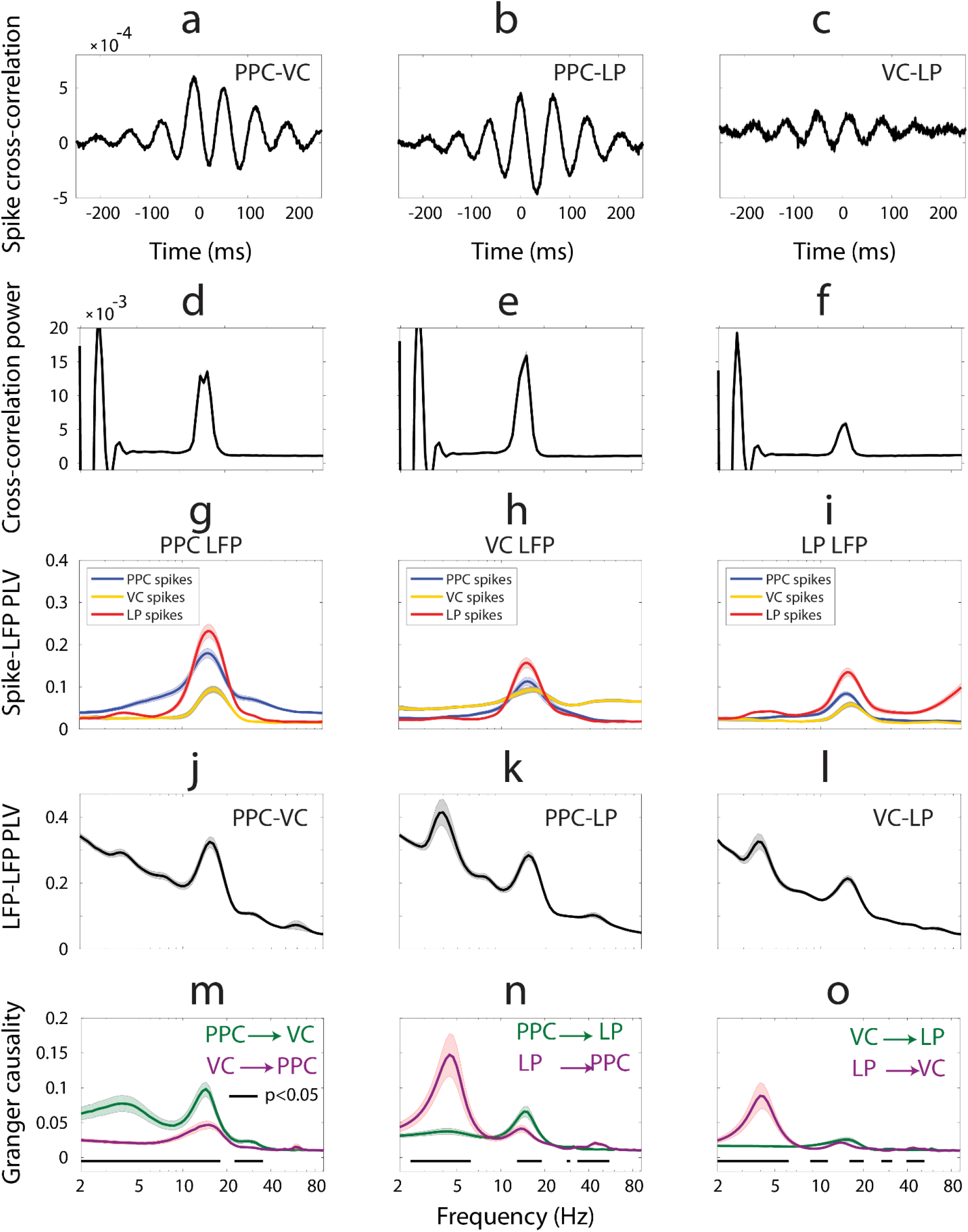
Synchronization between the regions and directed functional connectivity in thalamo-cortical network. Population average spike cross-correlation measured between PPC and VC **(a)**, PPC and LP **(b)**, and VC and LP **(c)**. The mean power spectrum of spike-correlations display a prominent peak in the alpha band (centered on ~15.23 Hz) for PPC-VC **(d)**, PPC-LP **(e)**, and VC-LP **(f)** spike crosscorrelations. **(g)** Phase locking between spikes from three ROIs and LFP from PPC region. **(h)** Phase locking between spikes from three ROIs and LFP from VC. **(i)** Phase locking between spikes from three ROIs and LFP from LP. **(j)** Phase-locking value (PLV) between LFP recorded from PPC and VC regions as a function offrequency indicates synchronization at alpha and weak theta frequency bands. **(k)** PPC and LP are synchronized in both theta and alpha-band frequency ranges. **(l)** Theta and alpha-band synchronization is evident between VC and LP regions. **(m)** Spectrally resolved Granger causality shows that the influence of PPC on VC (green) is larger than influence of VC on PPC (violet) in all examined frequencies including alpha band (p<0.05, t-test, indicated by horizontal line). **(n)** LP has a causal influence on PPC at theta frequency range (violet). In return, PPC has a causal influence on LP at alpha frequency band (green). (o) LP drives VC at theta frequency range (violet). VC and LP drive each other comparably (p>0.05, t-test) at alpha frequency range. All measures are shown as mean (thick line)±SEM(light background).

We found strong phase locking in the alpha range (peak frequency at 15.23 Hz) and weak synchronization in theta range (peak value at 3.75 Hz) between two cortical regions (Fig. 6 j). When considering the synchronization between a cortical region and LP, we found strong phase-locking at both theta and alpha frequency ranges (Fig. 6 k, l). The theta and alpha synchronization between PPC and LP was larger when compared to corresponding values for VC and LP. These results indicate that the two cortical regions synchronize in the alpha range, while the synchronization between a cortical region and LP develops in both alpha and theta frequency ranges as previously reported (Stitt et al., 2018). Further analysis by population average conditional Granger causality measure (cGC) showed that alpha synchronization between cortical regions was more directed from PPC to VC than the opposing direction (Fig. 6 m, see supplementary figure S1 for cGC measures for individual animals). The results of directed functional connectivity analysis also suggested that LP influences both cortical regions in the theta range, and on a smaller scale the PPC drives the LP in alpha range (Fig. 6 n, o). Our results did not indicate a preference in directionality of alpha oscillations between VC and LP (Fig. 6 o). Taking all together, these results show that alpha oscillations originate mainly in PPC and propagate to both VC and LP. On the other hand, the LP drives the other two regions in the theta frequency range.

### Measurements and modeling of tACS electric field in the ferret brain

We identified endogenous alpha oscillations that originated in PPC as a potential target for transcranial stimulation. To examine the effect of tACS on neurons, in our experiments we approximately matched the electrode current density of clinical tACS. The tACS current density is ~ 1 μA/mm^2^ in a return electrode of a typical dual-site clinical tACS setting where the current intensity is 2 mA peak for the return electrode (Ahn et al., 2019) We trimmed the clinical tACS electrodes to make smaller circular electrodes with radius of *r* = 5 mm to apply to the ferret head. To achieve a current density comparable to values used in human settings, we used current amplitudes up to 80 μA peak in our experiments.

We next examined what electric field strengths were generated inside the ferret brain when applying tACS currents with amplitudes ≤ 80 μA. To estimate the electric field magnitude in the ROIs and the entire ferret brain, we developed a computational model of the current flow in the ferret head (Fig. S6). The model included detailed anatomical structures, head-post, bone screws, dental cement, and surgical craniotomies based on MRI and CT scan data and tissue conductivities from the literature (Table S1). We validated the model adequacy by comparing its predictions to experimentally measured electric potentials for stimulation currents of 80 μA and 60 μA. The electric potentials were measured with the same implanted electrode arrays used for the electrophysiology recordings. The model prediction for the electric potentials was highly correlated with the experimental data (Fig. 7 a, b respectively, correlation coefficient r = 0.99, p < 0.0001 for both validations). The small error bars of each measurement in the plots stem from variation across the six sets of recordings corresponding to two recording blocks and three different reference electrode schemes, after re-referencing the data and excluding electrodes with poor signal quality (see Methods for details). The simulated distribution of the electric field magnitude in the three ROIs is summarized in Fig. 7 c. The estimated electric field magnitude was slightly higher for the VC region compared to PPC and LP, but most of the magnitudes across all ROIs fell into a relatively narrow range of 0.22–0.3 mV/mm. Figures 7 d, e show the surface-normal component of the electric field on the brain surface, corresponding to peaks and troughs of the tACS current cycle. The areas underneath the stimulation electrodes (indicated by large open circles on the left frontal and medial occipital regions) showed maximum field strength with directions flipped according to the stimulation wave polarity.

**Figure 7.**
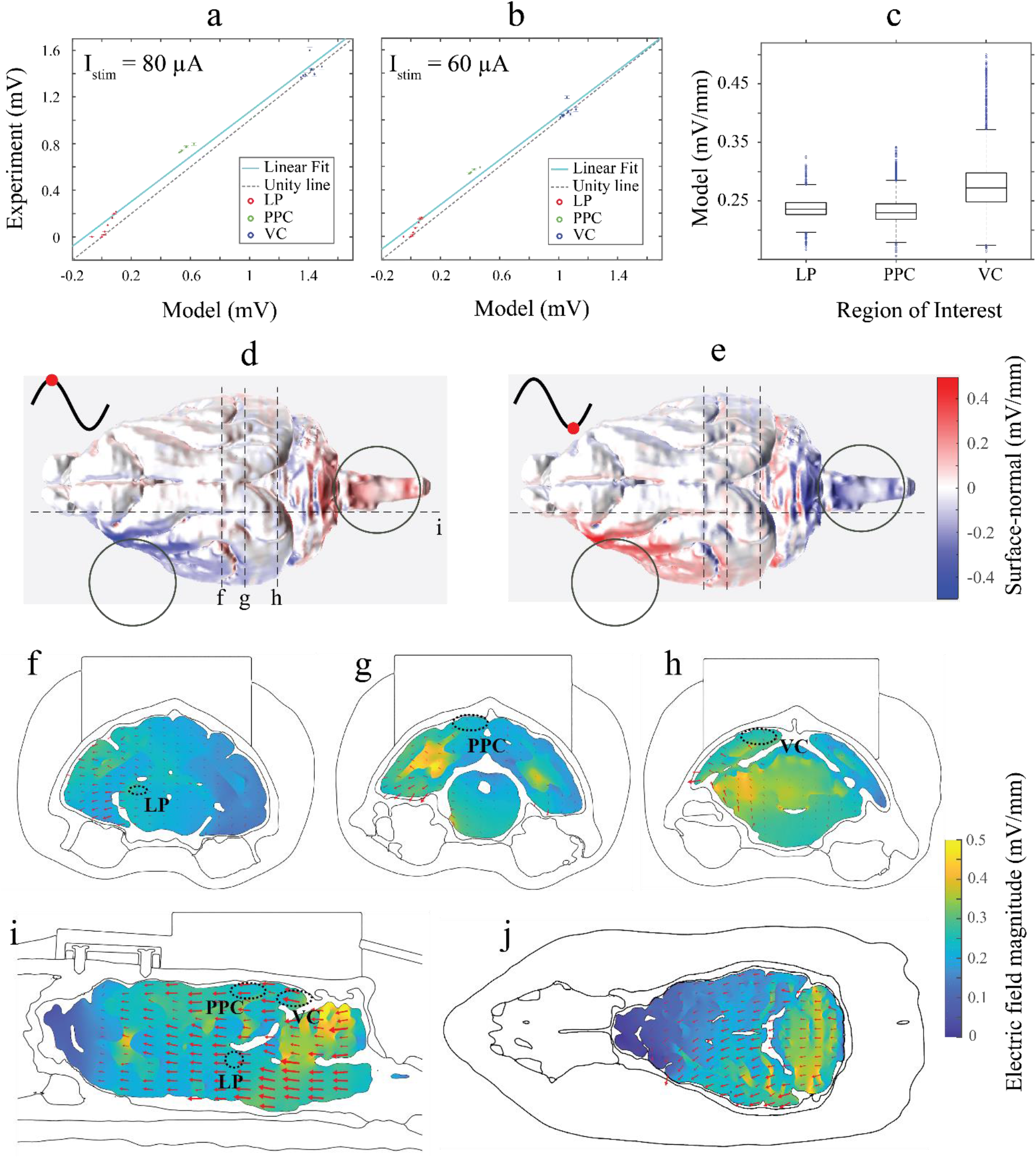
Measurement and modeling of tACS electric field in ferret brain. **(a, b)** Comparison between recorded and simulated electric potentials using stimulation current intensity of 80 μA and 60 μA in three recording sites: LP, PPC, and VC. The vertical error bars represent recording variability as standard deviation across the different reference schemes. Solid cyan line represents the linear regression of the data, which ideally should match the dashed black unity line. **(c)** Distribution of magnitude of threedimensional electric field vectors across three ROIs, predicted by model, indicating weak electric field of ~ 0.25 mV/mm. **(d)** Qualitative demonstration of spatial distribution of magnitude of surface-normal component of electric field on brain surface at 90 degrees phase of tACS wave (red dots on a cycle of sine wave) and **(e)** 270 degrees. The electric field magnitude is greatest in areas proximal to stimulation electrodes (open black circles in left frontal and midline occipital regions). The direction of the surface normal electric field flips according to the tACS phase. The electric field distribution along three coronal sections (including ROIs), one sagittal section (dashed lines in **d** and **e)** and a transversal section is displayed in **f-j**. The model prediction shows that the electric field magnitude is less than 0.5 mV/mm in all sections. The maximum electric field (bright yellow) occurs outside the ROIs. Arrows representing the direction of the electric field components are scaled by the electric field strength in the plane of the sections in **f-j** and demonstrate dominant posterior-anterior electric field direction.

Figures 7 f-i visualize the electric field strength and direction in deeper cortical and sub-cortical regions across coronal and sagittal sections including the implanted recording sites. As expected, the direction of the electric field was dominantly along the posterior–anterior axis as shown by red arrows in the coronal and sagittal sections.

These results indicate that neither the surface-normal nor the total magnitude of the tACS electric field exceeded 0.5 mV/mm in the brain. The modeling results also indicated that the arrangement of the tACS electrodes in our experiments did not necessarily result in maximum electric field in the PPC, VC, and LP regions where all cortical recordings were obtained. The electric field strength was in the range of values reported for humans and non-human primates in models and experiments (Huang et al., 2017; Opitz et al., 2016). These results support our choice of 80 μA as stimulation intensity resulting in comparable electric fields in human tACS settings at the sites of the implanted electrodes.

### Transcranial alternating current stimulation engages cortical alpha oscillations at the single neuron level

We found that synchronization between spikes and both local and large-scale oscillations was highest for alpha-band oscillations across all three animals. In addition we identified the PPC as the source of alpha oscillations that drives VC and LP. These results suggested alpha as a candidate target for tACS examinations. Further we identified that tACS with amplitude of 80 μA resulted in a weak electric field of < 0.5 mV/mm across the entire ferret brain. We next performed a comprehensive series of experiments where we applied tACS with different frequencies and amplitudes and simultaneously measured the spiking activity in PPC, VC and LP regions. Nine stimulation frequencies centered on each animal’s endogenous alpha frequency (*EAF*) (*f*_stim_ = [–4, –3, –2, –1, 0, 1, 2, 3, 4] + *EAF*) and 6 amplitudes (*I*_stim_ = [5, 10, 15, 20, 40, 80] *μ*A, peak values) were applied over multiple sessions per animal (*n*=16, 21, 14 sessions for animals 1 to 3 respectively). We hypothesized that if entrainment is the underlying mechanism of tACS effects, then the synchronization between endogenous oscillations and external tACS should increase as the tACS frequency approaches endogenous alpha frequency. This increased synchrony between an external periodic perturbation and an oscillator is a well-known phenomenon in dynamical systems. As the amplitude of the stimulation increases the range of frequencies over which a high synchrony is achieved also increases resulting in triangular Arnold tongue regions on twodimensional heat-maps constructed using stimulation amplitude and frequency values (Pikovsky et al., 2003).

We first analyzed the phase locking of all spikes to tACS waveform regardless of spike shapes in a collective dataset across three animals (see supplementary figures S2-S4 for analysis results for individual animals with results similar to those of collective dataset across animals). A triangular region centered on endogenous alpha frequency appeared for PPC units (Fig. 8 a top, darker shades of blue indicate higher PLV). We evaluated the modulation of spike phases using Rayleigh’s test for nonuniformity of circular data and found that the z-statistics also displays a triangular shape for PPC neurons (Fig. 8 a, second row). The percentage of PPC units showing significant phase modulation was low (<2%, Fig. 8 a, third row). The pattern of the corresponding firing rate map for PPC neurons was random without displaying a triangular shape. This confirmed that the examined tACS amplitudes were subthreshold and as expected for tACS, the stimulation did not modify the firing rate of the units (Fig. 8 a bottom). To further differentiate the cell-type-specific effect of tACS we extracted the synchronization maps for narrow- and broad-spiking neurons separately. We performed the same analysis on clustered units in PPC based on spike shapes: the narrow-spiking neurons (n=685) displayed darker triangular region when compared to their broad-spiking counterparts (n=864) (Fig. 8 b, c, top). The synchronization map between VC units and tACS (for collective or clustered spikes) did not show a regular Arnold tongue shape (Fig. 8 d-f top) as observed for PPC units. Instead, a weak modulation was observed for frequencies close to the endogenous alpha frequency, and became more evident for stimulation amplitudes above 20 μA and angled toward frequencies higher than endogenous alpha as indicated by dark region on top right corner in Fig. 8 d, e. We did not find high-synchrony regions for LP neurons (Fig. 8 g, top). As for PPC, the Rayleigh’s z-statistics in VC and LP (Fig. 8 d-g, second row) showed similar pattern to corresponding PLV maps (Fig. 8 d-g, top), and the percentage of units showing significant phase modulation was low (<2%, Fig 8 d-g, third row). Similar to PPC, tACS did not modulated the firing rate of individual units in VC or LP (Fig. 8 d-g, bottom row). Taken together these results show the synchronization map between single units and tACS that induced electric field of <0.5 mV/mm across the ferret brain. The synchronization region follows a triangular Arnold tongue shape, with phase locking values that are just slightly higher than their neighbor regions. The PPC units demonstrated Arnold tongue pattern indicative of their weak entrainment by tACS. The VC units demonstrated less regularly-shaped Arnold tongue and no regular pattern of synchrony was observed for LP neurons. Among narrow- and broad-spiking neurons, the former ones showed more developed Arnold tongue regions suggesting possibility of more entrainment by tACS.

**Figure 8.**
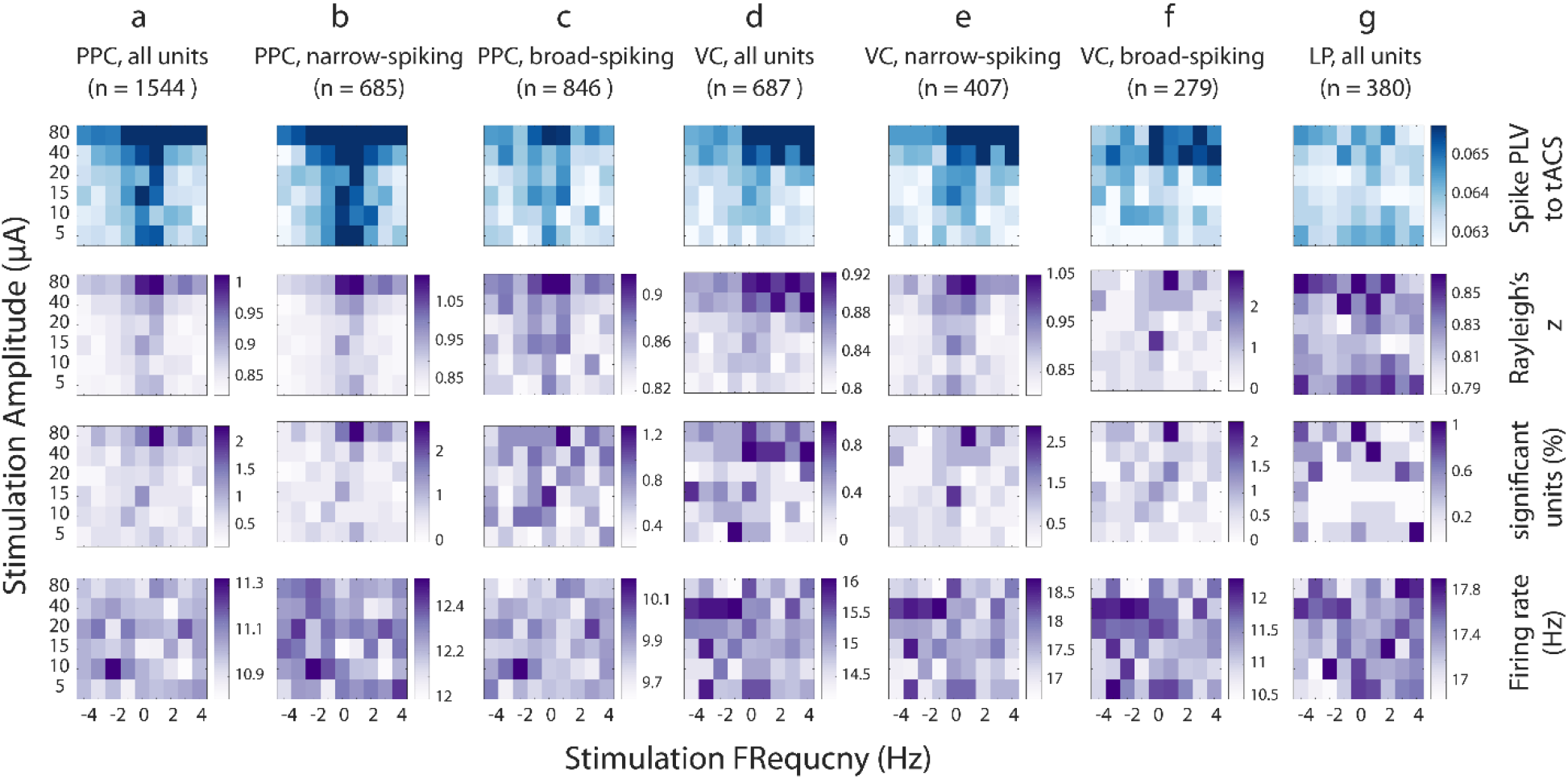
Transcranial alternating current stimulation engages cortical alpha oscillations in single neuron level. Synchronization maps (blue) and corresponding Rayleigh’s z-score (purple, second row), percentage of units with significant phase-locking value (purple, third row) and firing rate maps (purple, bottom row) as a function of stimulation parameters for PPC **(a-c)**, VC **(d-f)**, and LP **(g)**. The synchronization maps show phase locking between individual spikes and tACS wave as measured by phase-locking value (PLV) averaged across units. The horizontal axis indicates the distance (in Hz) from individual alpha frequency, and the vertical axis is the stimulation amplitude. (**a**, top two rows) tACS entrains PPC units as indicated by Arnold tongue centered at alpha frequency. (**b, c**, top two rows) Narrow-spiking units in PPC show stronger phase locking to tACS when compared to their broad-spiking counterparts. (**d-f**, top two rows) VC units are phase locked to tACS, but the synchronization maps are not well-developed triangular shape as observed for PPC. (**g**, top two rows) The synchronization map for LP units does not show a high-synchrony region. (**a-g**, third row) Only a small percent of units (<2%) show significant modulation of spike phases as measured by Rayleigh’s test. (**a-g**, bottom row) The random pattern of firing rate maps for all regions/unit types indicates that tACS did not modulate the firing rate of target neurons

### Endogenous alpha oscillations in a biophysical model of thalamo-cortical network and dynamics of synchronization by tACS

We adopted and connected two previously developed computational models of a cortical (Negahbani et al., 2018) and thalamic (Li et al., 2017) network for complementary investigation of neuronal mechanisms of tACS effect during endogenous alpha oscillations. The cortical model was composed of fast spiking inhibitory neurons (FS) and regular spiking excitatory pyramidal neurons (PY). The thalamic model included excitatory relay-mode thalamo-cortical cells (RTC), excitatory high-threshold bursting thalamic cells (HTC), inhibitory thalamic reticular cells (RE) and inhibitory thalamic interneurons (IN). Synaptic and gap junction connection patterns between neuronal populations followed previous modeling work (Fig 9 a). In the absence of stimulation, the network reproduced the major oscillatory patterns observed in experimental LFP recordings from cortex (PPC) and thalamus (LP) (Fig. 9 b). In the model, the simulated LFP (LFP) displayed dominant alpha (~ 10Hz) and weak theta-band (~3.9 Hz) oscillations in cortex and dominant theta and weak alpha in thalamus (Fig. 9 b, right, spectral graphs). We next stimulated PY neurons according to the same protocol we used in experimental investigation of entrainment by tACS. We first demonstrate how a sample tACS with frequency matching the endogenous alpha (10 Hz) and weak amplitude (1 pA) entrains individual neurons and the entire network. The network response to 10 Hz and 1 pA stimulation is shown in Fig 9 c, demonstrating the collective entrainment of different neuronal types to stimulation in terms of increased amplitude of cortical LFP compared to its amplitude without stimulation. The membrane potential of a randomly selected neuron from each neuron type in the network shows how individual FS and PY neurons adjust their spike times to tACS. We next applied a comprehensive set of tACS waveforms to cortical PY neurons with frequencies between 3 and 30 Hz and amplitudes between 1 and 10 pA, and computed the phase-locking of spikes to tACS for all neuron types (Fig. 9 d). We found Arnold tongue patterns (indicated by dark blue shades) centered on endogenous alpha frequency only for cortical neurons. Compared to cortical neurons, only a very weak entrainment was observed for thalamic neurons. These findings were in agreement with our experimental findings where tACS entrained cortical neurons in PPC, but not thalamic neurons in LP. Interestingly, we found that tACS entrained FS more than PY as indicated by wider and darker Arnold tongue pattern for FS. This behavior is in agreement with our experimental observations where narrow-spiking neurons of PPC showed larger phase-locking to tACS when compared to broad-spiking neurons. To clarify the neuronal response to tACS at different parts of the PLV heat-map, we further examined a PY neuron’s response to two tACS waveforms with equal amplitudes, but different frequencies: (i) A point on Arnold tongue (10 Hz, 8 pA), where PY neuron fires regularly after the peak of tACS, with a phase preference between 90 and 180 degrees as shown in phase histogram (Fig. 9 d top right), indicating a high synchronization by tACS. (ii) A point outside of Arnold tongue (15 Hz, 8 pA), where PY neuron fires at random phases of tACS, without a phase preference (Fig. 9 d bottom right), indicating a lack of synchronization by tACS. We next asked if the stimulation amplitudes we used in the model are a good estimate for experimental tACS amplitudes. The aim was to identify amplitude values in the model that resulted in subthreshold response in neurons, being a correct model for experimental tACS intensities. The extracted firing-rate heat-maps for all neuronal types are displayed in Fig. 9 d, bottom row. We observed a threshold value of stimulation amplitude (~ 2pA) for firing rate modulation in all neuron types. According to these results application of stimulation currents below 2 pA did not modulate firing rates (Fig. 9 d bottom row), while phase locked the cortical neurons (Fig. 9 d top row). We interpret *I*_stim_~2 pA as the maximum stimulation intensity in the model that does not modulate the firing rate of neurons while entraining their spiking as expected for tACS in experimental and clinical settings. Next we used the biophysical model of thalamo-cortical network to test if the frequency mismatch between dominant thalamic oscillations and alpha tACS was the reason for absence of Arnold tongues in thalamic records in our experiments. To answer this question we altered thalamic endogenous oscillations from being theta-dominated to alpha by increasing the DC drive current of HTC neurons from 0 pA to 50 pA and decreasing the drive current of RTC neurons from 100 pA to 75 pA. Then we stimulated the pyramidal neurons using the same tACS parameters as we used previously, and extracted the phase-locking maps. We found that both cortical and thalamic neurons were entrained by tACS and displayed Arnold tongues (Fig S5).

**Figure 9.**
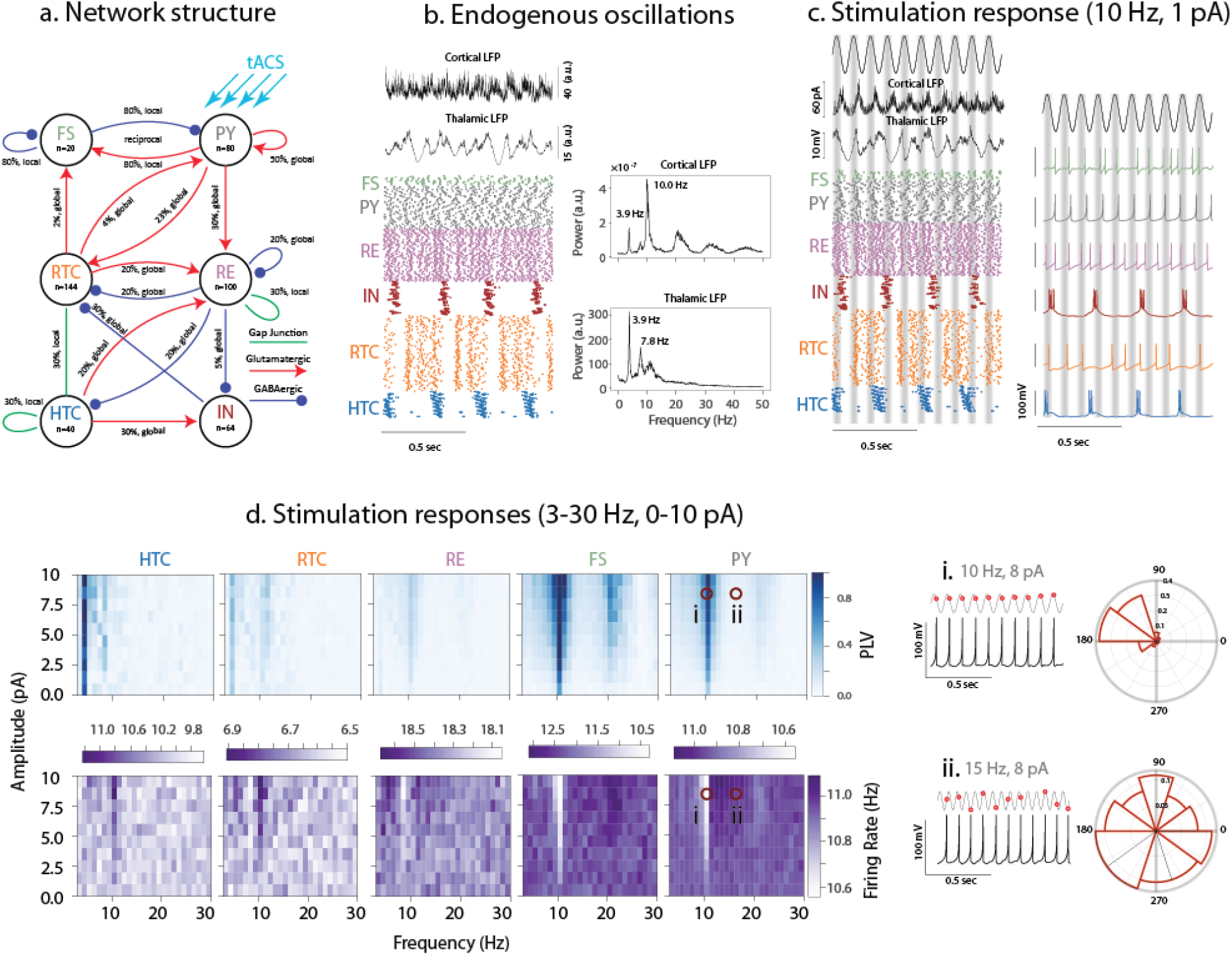
Endogenous alpha oscillations in a thalamo-cortical model and dynamics of synchronization by tACS. **(a)** The structure of the thalamocortical network with excitatory, inhibitory and gap-junction connections between neuronal populations. The cortical part includes fast-spiking inhibitory (FS) and regular-spiking pyramidal (PY) neurons. The thalamic part includes relay thalamo-cortical (RTC), reticular (RE), high-threshold bursting (HTC), and thalamic local inhibitory (IN) neurons. The tACS is applied only on pyramidal neurons (blue arrows). **(b)** Endogenous oscillations shown by (top) 1-sec cortical and thalamic local field potential (LFP) traces, and (bottom left) the raster plot with each dot representing the firing instant of a neuron. (Bottom right) The cortical LFP has a dominant spectral peak at alpha range (10 Hz), and a smaller peak at theta range (3.9 Hz). The thalamic LFP has a dominant theta peak (3.9 Hz), its harmonic (7.8 Hz), and a weak alpha peak (10 Hz). **(c)** Network response to a 10 Hz, 1 pA tACS. The cortical and thalamic LFP traces, and the raster plot showing firing of all neurons (left), and the membrane voltage traces of sample neuron from each cell type (right). **(d)** Network response to a group of stimuli with frequencies between 3 and 30 Hz, and amplitudes between 0 to 10 pA. Top left heat-maps: The color-coded phase locking value (PLV) as a function of stimulation frequency (horizontal axis), and stimulation amplitude (vertical axis) for each neuron type. Darker colors indicate higher synchronization between individual neurons and tACS with corresponding frequency and amplitude. The FS and PY neurons show triangular-shape high PLV regions centered on alpha (10 Hz). Bottom left heat-maps: The firing rate (FR) maps for each neuron type. The color at each point indicates the average firing rate (in Hz) across neurons in response to tACS with corresponding frequency (horizontal axis) and amplitude (vertical axis). Two points on PLV and FR maps are selected for further analysis: (i) A point with high PLV on Arnold tongue region with tACS parameters of 10 Hz and 8 pA. The tACS waveform and membrane potential of a sample PY neuron (top right) shows that the PY neuron fires regularly after the peak of the tACS wave (red circles). The phase distribution of tACS wave at firing instances of PY neuron shows a phase preference between 90 and 180 degrees. (ii) A point with low PLV outside of Arnold tongue region with tACS parameters of 15 Hz and 8 pA. The tACS waveform and membrane potential of a sample PY neuron (bottom right) shows that the PY neuron fires at random phases of tACS wave (red circles). The phase distribution of tACS wave at firing instances of PY neuron shows a phase distribution without any preference.

Together these modeling results provide support for our experimental data where we found stronger phase-locking in narrow spiking neurons when compared to broad-spiking ones in response to tACS.

## Discussion

### tACS entrains the spiking activity during endogenous alpha oscillations in awake head-fixed ferrets

We demonstrated the acute cellular-level effect of tACS on endogenous alpha oscillations and thereby provide experimental evidence for the previous predictions derived from computational modeling studies about entrainment of ongoing oscillations. We found that tACS entrained the spiking of individual cortical neurons but did not alter the neuronal firing rate during spontaneous alpha oscillations in absence of a task. We developed a finite element model of the ferret head, validated the model predictions with experimental measurement of tACS electric potential in the ferret brain, and tuned tACS amplitude to produce electric fields comparable to those reported in humans and nonhuman primates. We found how the phase-locking of spikes by tACS depended on stimulation frequency: matching the frequency of stimulation with endogenous alpha resulted in phase synchronization of spikes regardless of stimulation intensity. For tACS frequencies adjacent to endogenous alpha, higher stimulation intensities were required to achieve similar phase-locking between spikes and tACS. This behavior resulted in a triangular-shaped synchrony region when quantifying the spike-tACS phase-locking as a function of stimulation frequency and amplitude. The triangular region was centered on endogenous alpha frequency and is known as an Arnold tongue, as previously reported in computational modeling studies of tACS effect (Ali et al., 2013; Herrmann et al., 2016b; Lefebvre et al., 2017; Li et al., 2017; Negahbani et al., 2018). Despite being statistically significant in only a small fraction of neurons, the Arnold tongue was evident in PPC. The Arnold tongue was less developed and less clear in VC region, and was absent in LP region. Further we identified that effect of tACS was cell-type specific. The phase-locking between tACS and spikes was stronger for narrow-spiking neurons compared to their broad-spiking counterparts. This cell-type specificity was further supported by a computational model of thalamo-cortical network with ongoing alpha oscillations. In the model, the fast-spiking cortical neurons demonstrated higher phase-locking to tACS when compared to broad-spiking neurons.

### Weak electric fields (< 0.5 mV/mm) comparable to tACS field strength in humans and nonhuman primates can entrain neural spiking in the source of target oscillations

To examine the underlying mechanisms of how tACS modulates oscillatory interactions in the thalamo-cortical network, we here first delineated the functional interactions in the PPC-LP-VC network. We identified endogenous alpha oscillations in three regions based on three quantitative measures: LFP spectrum, phase-locking of spikes to population firing rate, and spike-EEG synchrony. Further analysis supported PPC as the source (or driver) of alpha oscillations in three-node network in resting awake ferrets. Despite this presence of alpha oscillations in all three recording regions and existing evidence for enhanced phase synchrony by weak electric fields when stimulation and endogenous frequencies are close (Alagapan et al., 2016; Neuling et al., 2013; Schmidt et al., 2014), we only found Arnold tongue pattern for PPC neurons.

Compared to PPC, the VC neurons showed less-developed triangular region when looking at PLV as a function of stimulation parameters. In addition the LP neurons did not demonstrate systematic phase locking to tACS. The magnitude of tACS electric field (model-driven value) was comparable in all three regions. Thus, we rule out differential field strength as the explanation. By PPC being the source of alpha oscillations, it may respond more strongly to tACS in the alpha frequency. In contradiction to recent reports of failure of electric fields less than 1mV/mm (resulting from commonly applied tACS intensities) in modulating the dynamics of neuronal circuits (Huang et al., 2017; Lafon et al., 2017; Vöröslakos et al., 2018), the identified PPC Arnold tongue in our work demonstrates how weak electric fields (<0.5 mV/mm) can engage individual neurons provided that the directionality of information flow and the source of target oscillations are considered in designing the stimulation protocol. Our results are the first *in vivo* demonstration of how a weaker electric field compared to the fields predicted and measured in human and nonhuman primate tACS studies (Opitz et al., 2016) can result in Arnold tongue regions on synchronization maps suggesting entrainment of neural populations.

### The head-fixed awake ferret model to study underlying mechanisms of alpha tACS effects

Modulation of spiking activity by externally applied subthreshold electric fields has been previously demonstrated in animal studies. Application of a subthreshold (< 10 mV/mm) 20 or 50 Hz oscillating electric field modulated the power and frequency of pharmacologically-induced gamma oscillations in CA3 pyramidal cells in rat brain slices (Deans et al., 2007b; Reato et al., 2010b). Fröhlich and McCormick showed how subthreshold oscillating electric field can enhance slow oscillations recorded from ferret visual cortex slices with effects demonstrated at field amplitudes as low as 0.25 mV/mm (Fröhlich and McCormick, 2010). Ozen et al. performed in vivo recordings from anesthetized and behaving rats while applying low intensity slow electric fields (<1.7 Hz) via electrodes placed on skull or dura, and reported reliable entrainment of neurons (Ozen et al., 2010). Ali et al. reported tACS-induced enhancement of slow endogenous oscillations in an anesthetized ferret with stimulation electrodes placed on skull (Ali et al., 2013). Despite the fact that these experimental reports present evidences for cellular-level effects of tACS, these studies share three common limitations. First, they have taken advantage of slow wave oscillations (~1 Hz, under anesthesia or *in vitro*) or fast gamma band oscillations (~ 30 Hz, pharmacologically induced, *in vitro*). None of these ongoing oscillations are the best model for the dominant ongoing oscillations in the awake brain. Second, using *in vitro* preparations or animal models with lissencephalic brains imposes limitations on the translation to human tACS. In addition, the way tACS electrodes were configured (bath application *in vitro*, and direct placement on dura or skull *in vivo*) does not match with standard practice for tACS electrode placement on the scalp in humans. Third, none of the abovementioned animal studies have explored the parameters of stimulation in a systematic way to present a comprehensive entrainment map as a function of stimulation frequency and amplitude. Our study overcomes these limitations by taking the advantage of using an awake head-fixed ferret animal model. Ferrets exhibit a gyrencephalic brain and display alpha oscillations in the awake state (Stitt et al., 2018), making them an appropriate choice for tACS animal studies beside nonhuman primates. In addition the placement of tACS rubber electrodes on the scalp with conductive gel also made our setting more comparable with commonly used strategies used in human tACS studies.

### Cell-specific effects of tACS

We identified two types of neurons based on their spike waveform shape in PPC and VC: narrow-spiking and broad-spiking neurons. With tACS, the phase of firing was modulated for both cell types, but the effects were more prominent for narrow-spiking neurons in PPC. In VC, despite the lack of a clear Arnold tongue structure, the narrow-spiking neurons also displayed stronger phase-locking to the stimulation waveform. The stronger coupling of narrow-spiking neurons to tACS was consistent with their behavior in absence of tACS. The narrow-spiking neurons showed stronger coupling to local (population firing rate) and large-scale (EEG) alpha oscillations when compared to broad-spiking neurons. The population ratio of narrow to broad-spiking neurons were 0.81 and 1.46 in PPC and VC, respectively. These ratios differ from the reported excitatory to inhibitory population ration of 0.25 in cortical regions of mammalian brain (Braitenberg and Schüz, 2013). Nevertheless, although layer V pyramidal neurons have been suggested to be the direct targets of tACS due to their elongated somato-dendritic axis, our findings suggested that tACS might also modulate the activity of other neuron types including fast-spiking interneurons as previously reported (Krause et al., 2013; Molaee-Ardekani et al., 2013; Moliadze et al., 2010; Radman et al., 2009; Reato et al., 2013). In our computational model of thalamo-cortical network that reproduced the key spectral features of our *in vivo* recordings in ferret PPC and LP, we applied tACS to only pyramidal neurons, and found that both pyramidal and fast-spiking neurons displayed phase-locking to tACS, with fast-spiking neurons showing slightly more phase-locking to tACS. This supports the notion that the response to tACS can be mediated via direct engagement or indirect network effects that is shaped by the endogenous activity of the cells and their role in the generation of the network activity patterns.

### Limitations

Like all scientific studies, our study has limitations. First, in tACS experiments, the phase locking values for the regions inside the Arnold tongues were small and only slightly higher than the corresponding values for adjacent regions. In addition only a small number of units (~ 2%) showed significant phase locking as tested by Rayleigh’s test. We propose that higher synchronization values might be measured if the electrophysiological recordings were acquired from the regions with higher tACS electric field strength as predicted by computational model of field distribution in ferret head. Second, we did not investigate stimulation frequencies in other frequency bands to examine the presence of additional Arnold tongues at the (sub)harmonics (Pikovsky et al., 2003). This was an experimental design choice based on time constraints for keeping ferrets comfortably head-fixed. We prioritized the collection of a large number of trials for the stimulation parameters that we examined. In agreement with previous modeling work of tACS (Ali et al., 2013; Li et al., 2018; Negahbani et al., 2018), we did observe an Arnold tongue at the harmonic of the endogenous alpha frequency in our computational model of the thalamocortical network. Third, we found comparable electric field magnitudes in cortex and thalamus for our stimulation montage. This demonstrates that the field strength on its own is not the main determinant of the effect on neuronal networks but rather that endogenous activity patterns and other factors shape the presence and magnitude of neuronal entrainment by tACS. Fourth, our electric field modeling results indicate that the electric field in cortical regions is largely in the posterior-anterior direction, which is perpendicular to the somato-dendritic axis of many pyramidal neurons. This might explain why broad-spiking neurons (potential pyramidal neurons) did not demonstrate stronger entrainment by tACS when compared to narrow-spiking neurons (potential non-pyramidal neurons) in our measurements. Ideally the stimulation electrodes should be placed so as to maximize the component of the electric field that is normal to ROI cortical surface to increase the field-induced polarization in pyramidal neurons in order to impose stronger phase-locking in target pyramidal neurons. The hardware-imposed constrains of implanted electrodes prevented us from placing tACS electrodes directly above the ROIs, which would have increased the electric field magnitude and component normal to the cortical surface. Fifth, our computational biophysical model of thalamo-cortical network was designed to include the required biophysical complexity to generate alpha oscillations but was not designed to capture all features of the experimental recordings, and was also not tuned to exhibit specific behavior to avoid artificial alignment of computational and biological results due to overfitting. Specifically, our model includes only a single cortical region, in contrast to our experimental recordings from two cortical area. In addition the endogenous alpha frequency is different between model (~10 Hz) and experiment (14-16 Hz in different animals).

In conclusion, we found that weak electric fields (< 0.5 mV/mm) comparable to the field strength measured in tACS experiments in humans and nonhuman primates with conventional stimulation current intensities can entrain neuronal activity. In addition our findings provide the first in vivo evidences to support the model-driven predictions about how tACS entrains ongoing neuronal oscillations as demonstrated by Arnold tongue pattern. We show how important is it to consider the endogenous activity patterns, their directionality, and spatial source of target oscillations to be able to entrain the neurons, as tACS entrained PPC but not VC and LP neurons in our experiments. Our findings re-emphasize the importance of animal models in tACS studies by providing evidence for effectiveness of weak electric fields in modulating the network oscillations via entraining the activity of neuronal populations.

## Acknowledgement

The authors would like to thank Angel Huang and Charles Zhou for comments on the manuscript, and Connie S. Scoggins, John M. Bernabei, Dr. Alexandra Badea, Neerav Goswami, Dr. Luis Gomez, and Dr. Lari Koponen for advice and technical assistance on the electric field modeling. This work was supported by NIH grant R01MH111889. The content of current research is solely the responsibility of the authors and does not necessarily represent the official views of the National Institutes of Health.

## Author Contribution

EN, IMS, and FF designed the experiments. IMS and EN performed the implant surgeries. IMS performed the experiments and collected the electrophysiology data. EN collected the potential and electric field measurement data. EN and IMS analyzed the experimental results. EN performed the histology experiments. SRS interpreted the histological data and contributed text and figure to the manuscript. MD performed biophysical modeling of thalamo-crotical network, analyzed the results, and contributed text and figures to the manuscript. TTD, ACH, MD and AVP created and analyzed the finite element model of the electric field in a ferret head. TTD, MD, and AVP analyzed the electric potential recordings and contributed text to the manuscript. EN and FF wrote the manuscript. All authors reviewed the manuscript.

## STAR Methods

### Animals

Three adult spayed female ferrets (*Mustelaputoriousfuro*, 4 months old at the beginning of the experiment) were used in this study. All animal procedures were performed in compliance with the National Institute of Health guide for the care and use of laboratory animals (NIH publication No. 8023, revised 1987) and the United States Department of Agriculture, and were approved by the Institutional Animal Care and Use Committee of the University of North Carolina at Chapel Hill.

### Head-post and electrode implant surgery

The initial induction of anesthesia was performed with intramuscular injection of ketamine/xylazine (30 mg/kg of ketamine, 1-2 mg/kg of zylazine). After confirming the loss of paw pinch reflex, animals were intubated for isoflurane (0.5-2% in 100% oxygen) delivery via mechanical ventilation. The physiological parameters including electroencephalogram, partial oxygen concentration, end-tidal CO2, and rectal temperature were continuously monitored throughout the surgical procedure to maintain the animal in a stable state. All surgical procedures were performed under sterile conditions. The skull was fixed to a stereotactic frame using a mouth-piece and ear bars to enable accurate identification of target regions and electrode implantations. A custom-designed stainless steel head-post was first secured to the anterior part of the exposed skull via four stainless steel bone screws. A craniotomy was performed over the left hemisphere of PPC to implant Microelectrode arrays in PPC and LP regions. Another craniotomy was performed over the left hemisphere of VC region. The dura and then pia were removed before lowering the LP microelectrode array (2 × 8 tungsten electrodes, 35 μm diameter, 9 mm length, 250 μm spacing; Microprobes for life science). The second microelectrode array (2 × 8 or 2 × 16 tungsten electrodes, 35 μm diameter, 9 mm length, 200 μm spacing; Innovative Neurophysiology) was lowered to deep cortical layers in lateral gyrus in PPC region. The third microelectrode array (2 × 8 tungsten electrodes, 35 μm diameter, 9 mm length, 200 μm spacing; Innovative Neurophysiology) was implanted in VC area after removal of dura and pia layers. The reference electrode was directly adjacent to recording electrodes for all three microelectrodes. Each microelectrode array had a silver wire for ground connection. Three bone screws on the other hemisphere were used as ground for each microelectrode array. The EEG bone screws were placed on both sides of PPC craniotomy region. Microelectrode arrays were fixed in place with dental cement. After hardening of dental cement, the muscle and the skin around the incision were sutured together. Animals were administrated preventative analgesics and antibiotics for one week after surgery, while recovering in their home cage for at least one week prior to recordings.

### Transcranial electric current stimulation procedure

Two circular tACS electrodes (r = 5 mm) were prepared by trimming the conventional tACS electrodes. The electrodes were applied to ferret head using Ten20 conductive EEG paste over the left frontal area (directly above left eye) and central posterior area (over the neck). The digital tACS waveforms with desired timing and frequency were first generated using a Matlab script, then converted to analogue signal using a National Instruments DAQ device. The analogue outputs were connected to an A395 linear stimulation isolator (World Precision Instruments) to produce the final tACS currents that were delivered to animal’s head via tACS electrodes. For each stimulation session, the head-fixed animal received 54 stimulation block, each 90 sec with 10 sec inter-stimulus interval. Each stimulation block was randomly selected from a pool of 54 different combination of stimulation amplitudes (5, 10, 15, 20, 40, 80 μA) and frequencies (alpha-4, alpha-3, …, alpha, …, alpha+3, alpha+4 Hz). The stimulation and simultaneous electrophysiological measurements were performed in a room with dim lights while animal was awake and not engaged in any particular task.

### Electric potential measurements

To validate the electric field model of tACS in the ferret brain, we collected intracranial measurements of electric potentials from the electrode arrays in PPC, VC, and LP for one animal across three recording sessions. The potentials of the electrodes in all three ROIs were referenced to a single electrode potential in the proximity of a particular ROI that was changed for each recording session (for simplicity all sessions were subsequently re-referenced to the first electrode channel of the LP electrode array). During these in-vivo measurements, sinusoidal electric current was injected through tACS scalp electrodes using a fixed current amplitude of 60 or 80 μA. Four blocks of measurements were recorded in an alternating pattern (60, 80, 60, 80 μA); other parameters were kept constant: frequency = 14 Hz, measurement trial duration = 3 min, interval between trials = 24 s). Thus, for each tACS current amplitude there were six recording sets (two blocks and three reference schemes). The electrode potentials were recorded with a sampling frequency of 333.33 Hz, and were passed through a digital band-pass filter (order: 263, generalized equiripple filter with passband of 12-16 Hz, stopband cutoff frequencies = 8, 20 Hz; stopband gain = 90 dB; passband gain = 1 dB). To extract reliably the signal amplitude for each electrode, the recorded potentials were fitted to a reference sinewave using the MATLAB function fit. The reference waveform was derived by fitting a sine function to the potential waveforms from the electrode array (LP, PPC or VC) with largest signal per recording set. The sine frequency and phase estimates for each available array electrode (r^2^ > 0.997) were averaged to define the reference waveform. The amplitude of the reference sinewave was then fitted to the potential from each electrode in each recording set, and data from noisy electrodes (r^2^ < 0.75) were discarded. The standard deviation across the recording sets was computed for the remaining fitted amplitudes within each electrode, and the MATLAB isoutlier function (with default settings) was used to exclude electrodes with large standard deviation, and therefore poor reproducibility. The amplitude estimates were averaged within the remaining electrodes. Out of 9 nominally connected electrodes per array, 8, 6, and 8 electrodes survived these quality checks for the LP, PPC, and VC arrays, respectively.

### Computational modeling of electric field distribution in ferret head

We developed a finite element model for a ferret head to study the tACS electric field distribution in the brain. The modeling pipeline is comprised of five steps: image acquisition, image registration, image segmentation and implant model, complete model meshing, and electric field calculation (see supplementary Fig. S4). CT and MRI images of an in vivo ferret head without any surgical alterations were collected and manually co-registered. These images were segmented into six tissue regions with different electrical properties: white matter, grey matter, cerebrospinal fluid, nasal cavity (air), skull, and scalp. The image segmentation was modified to include craniotomy tissue alterations, such as implanted hardware and insulating materials based on CAD models. The three dimensional model was meshed into finite elements. Using previously reported values (Huang et al., 2017; Kumar et al., 2014; Steels, 2008) a tissue electric conductivity was assigned to each element of the mesh (Table S1). Finally, the mesh was solved using the finite element method to calculate the electric potentials and electric field.

#### Image acquisition

In-vivo magnetic resonance imaging (MRI) of an animal without any surgical alterations was acquired in a Bruker 9.4 T scanner (Bruker AVANCE, Billerica, MA). The MR scan was collected using a RARE-sequence, with TR = 2000 ms, TE = 9.610 ms, voxels = isotropic 0.2 mm3, FOV = 42×42×36 mm, 210 sagittal slices. Ex-vivo computed tomography (CT) imaging of the same animal was acquired on a eXplore CT 120 (GE Healthcare) scanner. The CT scan was collected with isotropic 0.0995 mm^3^ voxels and matrix dimension = 430×363×883. Ex-vivo CT imaging of a different animal with craniotomy was acquired on a eXplore speCZT (GE Healthcare) scanner. The CT scan was collected with isotropic 0.0995 mm^3^ voxels and matrix dimension = 430×363×883.

#### Tissue Segmentation

Ferret head model generation began with manual image co-registration of the MRI and CT scans using 3D Slicer (Fedorov et al., 2012). To correct for bias field inhomogeneity, N4ITKBiasFieldCorrection (Tustison et al., 2010), a 3D Slicer tool, was applied to the MRI prior to co-registration. This co-registration is able to combine the soft tissues and air seen in the MRI with the hard tissue (skull) seen in the CT. The co-registered images were manually segmented using 3D Slicer into six different regions: white and grey matter, cerebrospinal fluid, nasal cavity (air), skull, and scalp. Further, the three brain ROIs (LP, PPC, and VC) were defined, including both white and grey matter. To avoid problems during meshing of smoothed surfaces derived from the segmentation, we up-sampled the segmentation from 430×363×883 isotropic voxels (0.0995 mm^3^ each) to 538×454× 1104 isotropic voxels (0.0796 mm^3^ each). Smoothing and gap filling of the segmentation were performed by utilizing the interactive segmentation software SimpleWare ScanIP (SIMPLEWARE Ltd., Exeter, UK).

#### Finite element meshing

All hardware affixations were reconstructed using SolidWorks from the registered CT data set, positioned in the model, and converted into segmented regions with the aid of photos taken after surgery, unless specified otherwise. The model represented a post-surgery ferret head with craniotomy, implanted hardware components, and insulating materials used in the experimental setup. Images of the electrodes were used in the reconstruction and positioning of realistically-shaped tACS electrodes on the model ferret head. The head post, head post bone screws, dental cement, and acrylic glue were represented in the model. The recording electrode arrays were not modeled as they were too small and had a high electrical impedance. The head post was reconstructed and positioned based on post-surgery CT imaging and photos, replacing skull voxels in that region. The head post bone screws were reconstructed using manufacturer datasheets and positioned into the screw holes of the head post, replacing skull voxels in that region. The acrylic glue was reconstructed with a rectangular prism shape that was modified to wrap around the head post and the head of the bone screws by subtracting skull, head post, and bone screw voxels from it. The dental cement filling the craniotomy was reconstructed by merging a cylinder and a rectangular prism shape and wrapping the composite around the head post and dental cement. The shape replaced scalp voxels in that region and was subtracted from the skull, head post, and acrylic glue voxels. To prevent overlapping, all tissues and materials were subtracted from each other and combined together from the inside out. The ScanFE module in ScanIP was used to convert the segmented images of both head models to finite element meshes. The resulting mesh consisted of 4.2 million tetrahedrons with 0.74 million nodes.

#### Electric potential and field computation

The finite element mesh was imported into COMSOL Multiphysics 4.3a (COMSOL Inc., Burlington, MA, USA). A set of isotropic conductivity values were assigned to the mesh as listed in Table S1. The top surface of the occipital stimulation pad was injected with a fixed current of 60 or 80 μA, while the top surface of the frontal stimulation pad was assigned as ground. To calculate the electric potential and field distribution, the quasi-static Laplace equation

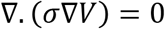

was solved with appropriate boundary conditions using the preconditioned conjugate solver and a relative tolerance of 10^−6^, where V and σ represent the electric potential and electric conductivity, respectively. Using the quasi-static approximation makes it possible to scale the stimulation current linearly to achieve the desired electric potential or electric field. The electric potentials were extracted from specific coordinate points in the model corresponding to the ideal ROI electrode locations, as the ROI electrodes were assumed to be point electrodes.

### Verifying electrode positions with histology

The animals were euthanized with an overdose of ketamine/xylazine after reaching the scientific endpoint and perfused immediately with 4% phosphate buffered paraformaldehyde. After removing the brain from the skull, it was post-fixed overnight in the same solution, cryoprotected in 30% phosphate buffered sucrose solution, shock frozen in dry ice and cut into 50 μm thick sections with a with a cryostat (CM3050S, Leica Microsystems). Sections were separated into two series and stained for cells (Nissl) or cytochrome oxidase. Imaging was conducted with an Aperio VERSA bright-field slide scanner at 10x magnification. The electrode arrays were located in the sections by electrode tracks, tissue damage or loss and atlas based reconstructed by comparing and documenting the sections to ferret atlas plates (Radtke-Schuller, 2018).

### Electrophysiological data analysis

Spectral domain analysis: We used wavelet transform for spectral decomposition. The wavelets (*w*(*t*, *f*_0_)) have Gaussian shape both in time and frequency:

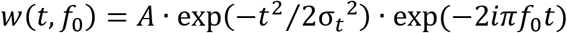

where *f*_0_ is the central frequency, 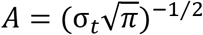 is a normalization factor to make the wavelet energy equal to 1, and σ*_t_* is the time domain standard deviation that relates to frequency domain standard deviation (σ*_f_*) as

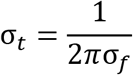

The frequency domain standard deviation is defined as a constant depending on central frequency of the wavelet as σ*_f_* = *f*_0_/7 (Tallon-Baudry et al., 1996). The local field potential or EEG signal was then convolved with Morlet’s wavelet to produce the complex-value analytical signal *X*(*t*, *f*_0_) at each frequency of interest. We used a family of 100 Morlet wavelets with central frequencies logarithmically spaced between 2 and 100 Hz. Then the power spectrum of the signal was computed by squaring the absolute value of analytical signal. For a given region of interest, session and animal, the power spectrum traces corresponding to all channels were plotted collectively to visually identify and exclude noisy channels. For each animal and for each region of interest we computed the average power spectrum as the mean spectrum across channels and then across sessions.

Spike-extraction, sorting and clustering: LFP signals were first high-pass filtered (300 Hz, 4^th^ order Butterworth filter). We determined the filter order by comparing the spectral contents of filtered signal using different filter orders to fully suppress the stimulation. The spiking threshold was set to negative four times standard deviation of the high-passed filtered signal, to extract spike times and spike waveforms. The extracted spikes were automatically sorted based on a template matching method implemented in Kilosort (Pachitariu et al., 2016), followed by an automatic post-hoc merging, and finally a manual curation by merging the automatically assigned spike groups. Next we used spike half-width to perform k-means clustering of neurons to two putative subgroups of narrow- and broad-spiking neurons (Barthó et al., 2004). Increasing the number of clusters did not improve the clustering performance as measured by within cluster sums of point to centroid distances. Other spike shape features (trough-to-peak time, peak value of normalized spikes, upward slope, spike energy, absolute peak-to-trough ratio, repolarization area, spike minimum amplitude, spike maximum amplitude) did not have multi-modal distributions and were not used in spike clustering.

Phase locking value (PLV): The consistency of phase differences between two signals is indicative of an existing phase synchrony and can be measured by PLV (Lachaux et al., 1999). We computed the PLV to quantify the phase synchrony between two LFP signals or the phase synchrony between individual spikes and tACS waveform. According to the instantaneous phase of tACS waveform at a specific carrier frequency (*f_c_*), we assigned a phase to every spike of a single unit, and built a phase distribution for each unit. Then we computed PLV of each unit with *N* spikes at a carrier frequency of *f_c_* defined by the following formula:

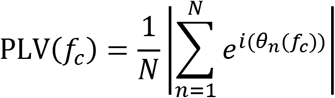

where *θ_n_*(*f_c_*) is the instantaneous phase at frequency f_c_ assigned to a spike. We used fixed number of *N* = 200 randomly selected spikes per unit, computed PLV, and repeated for each unit 200 times, with the mean PLV across all permutations as the final result. We assessed the significance of modulation of spike phases by tACS using Rayleigh’s test of uniformity, controlled for number of spikes (Zar, 1999). The PLV between spikes and LFP was computed in a similar way with the exception that spike phase were determined using LFP phases, and a fixed number of 1000 spikes were used to extract the PLV. The PLV between two LFP signals was computed in a similar way as used for computation PLV for spike-tACS, with the exception that in this case *θ_n_*(*f_c_*) was the difference between instantaneous phases of two LFP signals.

Directed functional connectivity: The Wiener-Granger causality algorithm (Geweke, 1982; Geweke, 1984; Granger, 1969) infers the influence of a process *y* on a process *x* by building two autoregressive models: a full model where previous values of both x and *y* processes are used to estimate the current value of *x*, and a reduced model where the prediction of current *x* value is made solely based on previous *x* values. The algorithm then compares corresponding residuals from two models and infers the directionality from *y* to *x* if the full model results in less residual error. We used MVGC toolbox (Barnett and Seth, 2014) to apply conditional form of Wiener-Granger causality analysis in frequency domain to study the directionality of different oscillatory frequencies in a network of three nodes at PPC, VC and LP. The MVGC toolbox models the observed data using a vector autoregressive (VAR) approach without assuming a linear scheme for observed real data. The process starts with estimating the model order in MVGC toolbox using Akaike or Bayesian information criteria or cross validation. The next step is to estimate the parameters of VAR model for both full model (considering the effects of both time series) and reduced model (considering only one time-series). Finally the Granger causality measure will be estimated based on the estimators of residuals of covariance matrices. We performed directionality analysis on LFP time series down-sampled to 200 Hz, then windowed into segments of 10 seconds length (with %75 overlap). LFP time series were collected from 10 randomly selected channels from each region, the directionality analysis was performed on all possible pairs, then averaged across to produce final results for PPC-VC, PPC-LP, and VC-LP pairs. We used Akaike method for model order estimation with maximum allowed model order of 20.

Spike triggered population firing rate and spike-triggered EEG: We computed the population firing rate function for each region of interest using spike times collected from that region via multiple channels of implanted microelectrode array. For each region, the spike times from all channels were pooled together, then sorted, then a binary time-series were constructed based on sorted collective spike times with assigned value of one at time of spikes occurrence, and zero otherwise. We next convolved the binary time-series with a Gaussian kernel with standard deviation of *c* = 4.6 ms and kernel width of 30 ms. We constructed the spike-triggered population firing rate and spike-triggered EEG by extracting a 1-sec window of population firing rate or EEG around each spike, then averaging across spikes.

### Computational modeling of thalamo-cortical network

We adopted our previously developed computational models of cortex (Negahbani et al., 2018) and thalamus (Li et al., 2017) and connected them in a biologically plausible way (Izhikevich and Edelman, 2008) to form the unified thalamo-cortical (TC) model. In short, cortical part includes 80 pyramidal (PY) and 20 fast spiking inhibitory (FS) neurons, spatially arranged on a line. Connections within PY neurons is global with connection probability of 0.5. Each FS was connected to 10 adjacent FS neurons with probability of 0.8. The reciprocal PY-FS connection was local where every FS was randomly connected to 0.8 of 32 surrounding PY. A biophysical synaptic model based on α-amino-3hydroxy-5-isoxazolepropionic acid (AMPA) or γ-aminobutyric acid type A (GABA_A_) mediated synaptic currents was used to connect cell populations. Individual PY and FS cells were modelled using Izhikevich formalism for point neurons (Izhikevich, 2007) with same parameter values reported previously (Negahbani et al., 2018). The thalamic network included 144 relay-model thalamic cells (RTC), 100 reticular inhibitory neurons (RE), 49 high-threshold bursting thalamic cells (HTC) and 64 local interneurons (IN), all placed in a two dimensional grid. All chemical synaptic connections were global in thalamic model with connection probability of 0.3 for HTC→·IN, ΓN→·RTC, and 0.2 for HTC→·RE, RTC→·RE, RE→RTC, RE→·RTC and RE→·RE. All gap junction connections were local, with connection probability of 0.3. All thalamic neurons were modelled using Hodgkin-Huxley formalism of point neurons, connected with glutamatergic (mediated by both AMPA and NMDA receptors) and GABAergic (mediated by GABA_A_ receptors) synaptic currents (with implemented short-term synaptic depression), or via gap junctions, with parameter values described previously (Li et al., 2017). The synthetic local field potential (LFP) for cortex was modelled as the average sum of the absolute values of excitatory and inhibitory currents entering PY neurons. The thalamic LFP was obtained from average membrane potential of neurons. The tACS was modelled as a current injection into PY neurons. The computational modeling of thalamo-cortical network was implemented using Brian2 simulator in Python.

### Statistical analysis

The comparison of means for two populations were performed using independent t-test with significance levels at 0.05 or 0.01. The significance of phase-locking values were assessed by Rayleigh test for non-uniformity of circular distribution of phases.

### Data and software availability

Electrophysiological data and MATLAB code that was used to perform analysis and visualize the results are available upon request from the corresponding author.

## Supplemental Information Titles and Legends

**Figure S1.**
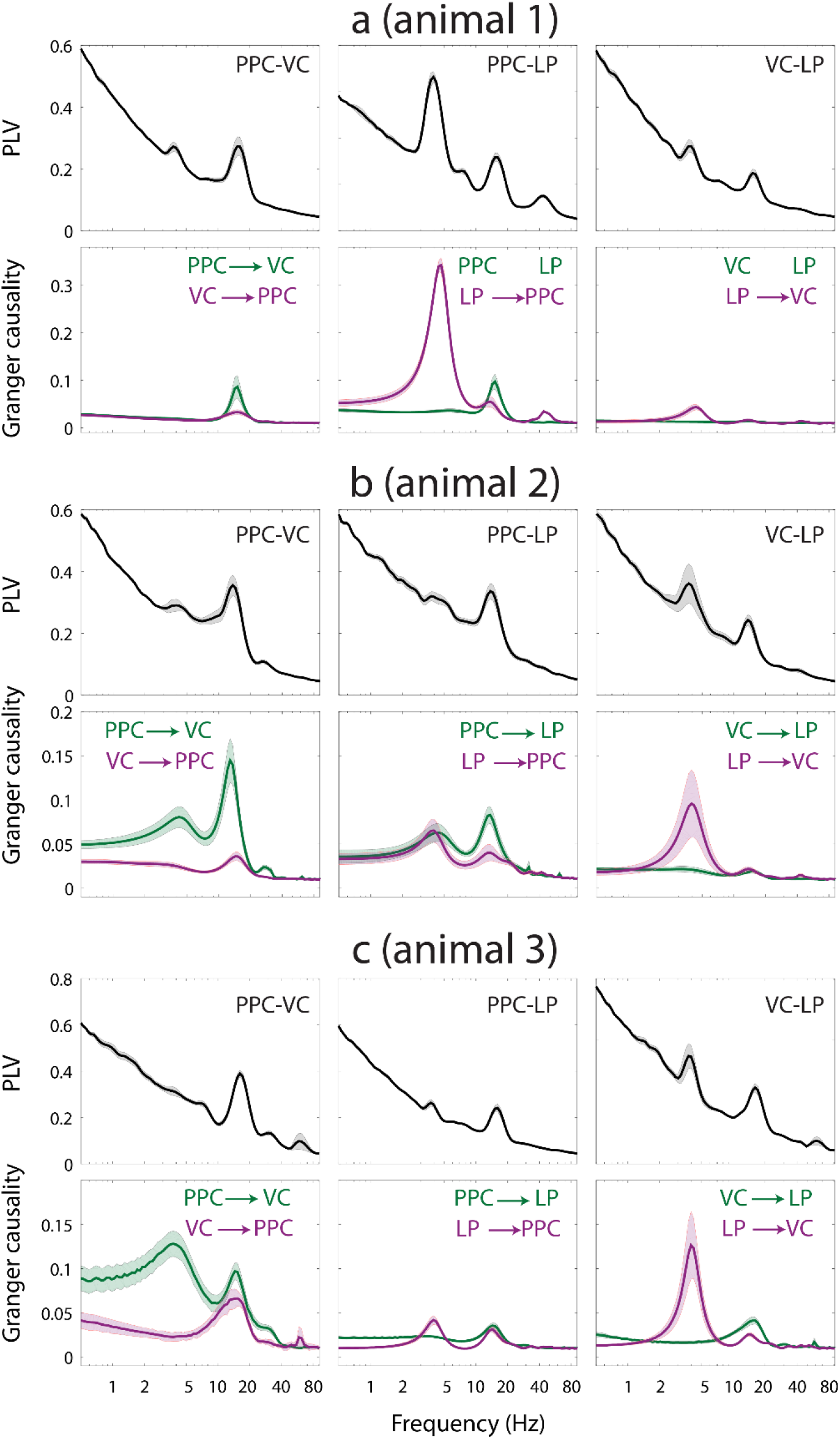
Synchronization between regions and directionality analysis for individual animals. For each animals, the top row shows the frequency dependent phase-locking values (PLV) and the bottom row displays the spectrally resolved conditional Granger causality results. Related to Figure 6.

**Figure S2.**
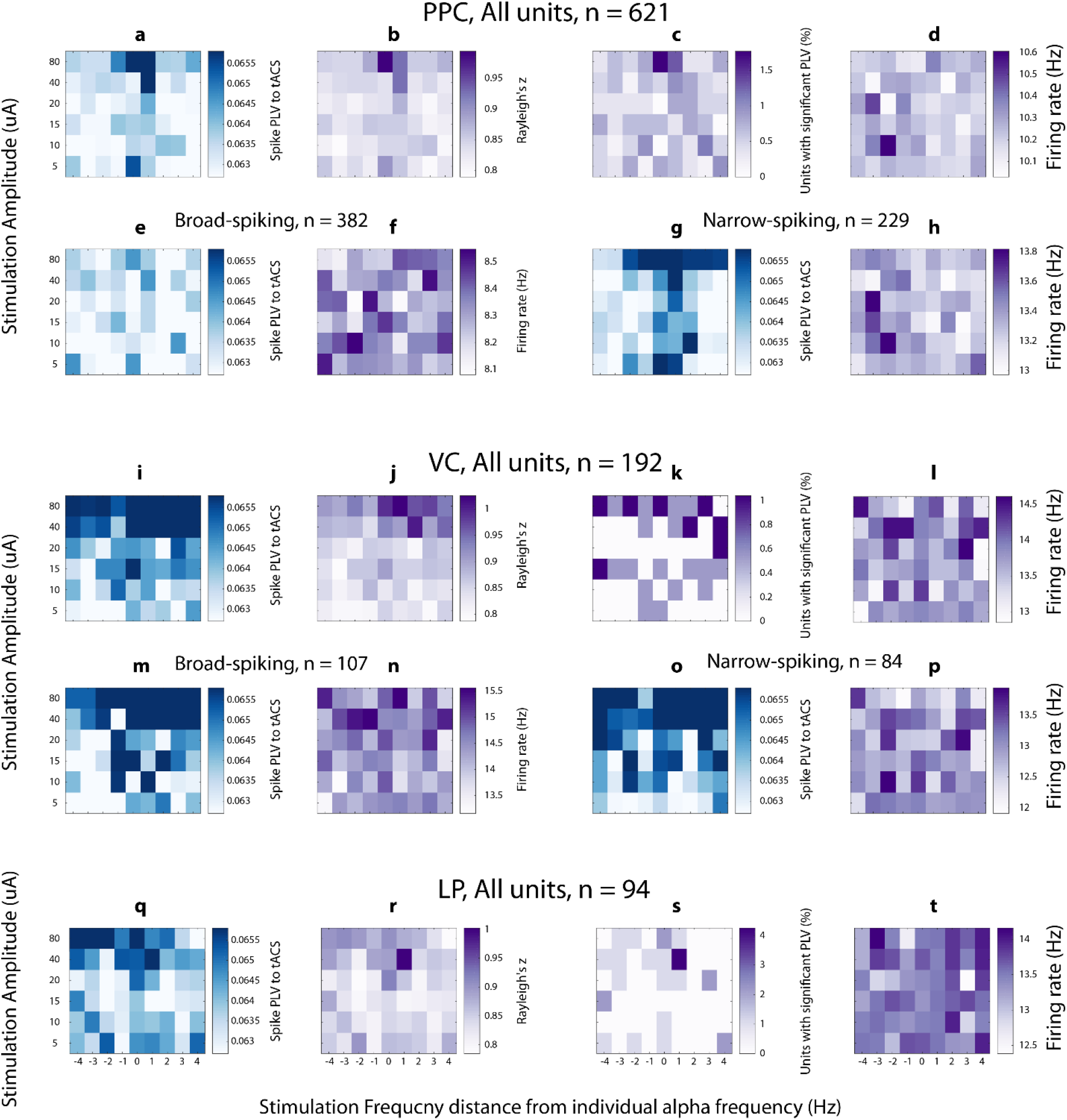
Synchronization and firing rate maps for animal 1. **(a)** Average phase locking of PPC spikes (averaged across 621 units) to tACS phase as measured by phase-locking value (PLV) as a function of stimulation frequency (horizontal axis) and stimulation amplitude (vertical axis). **(b)** Average Rayleigh’s z-score (across all PPC units) as a function of stimulation parameters. **(c)** Percentage of PPC units with significant PLV as indicated by Rayleigh’s test. **(d)** Average firing rate of all PPC units. **(e)** Average PLV of broad-spiking PPC units (n=382). **(f)** Average firing rate of broad-spiking PPC units. **(g)** Average PLV of narrow-spiking PPC units (n=229). **(h)** Average firing rate of narrow-spiking PPC units. **(i-p)** Corresponding measures of **(a-h)** for VC. **(q-t)** Corresponding measures of **(a-d)** for LP. The horizontal axis shows the distance from endogenous alpha frequency in Hz. Related to Figure 8.

**Figure S3.**
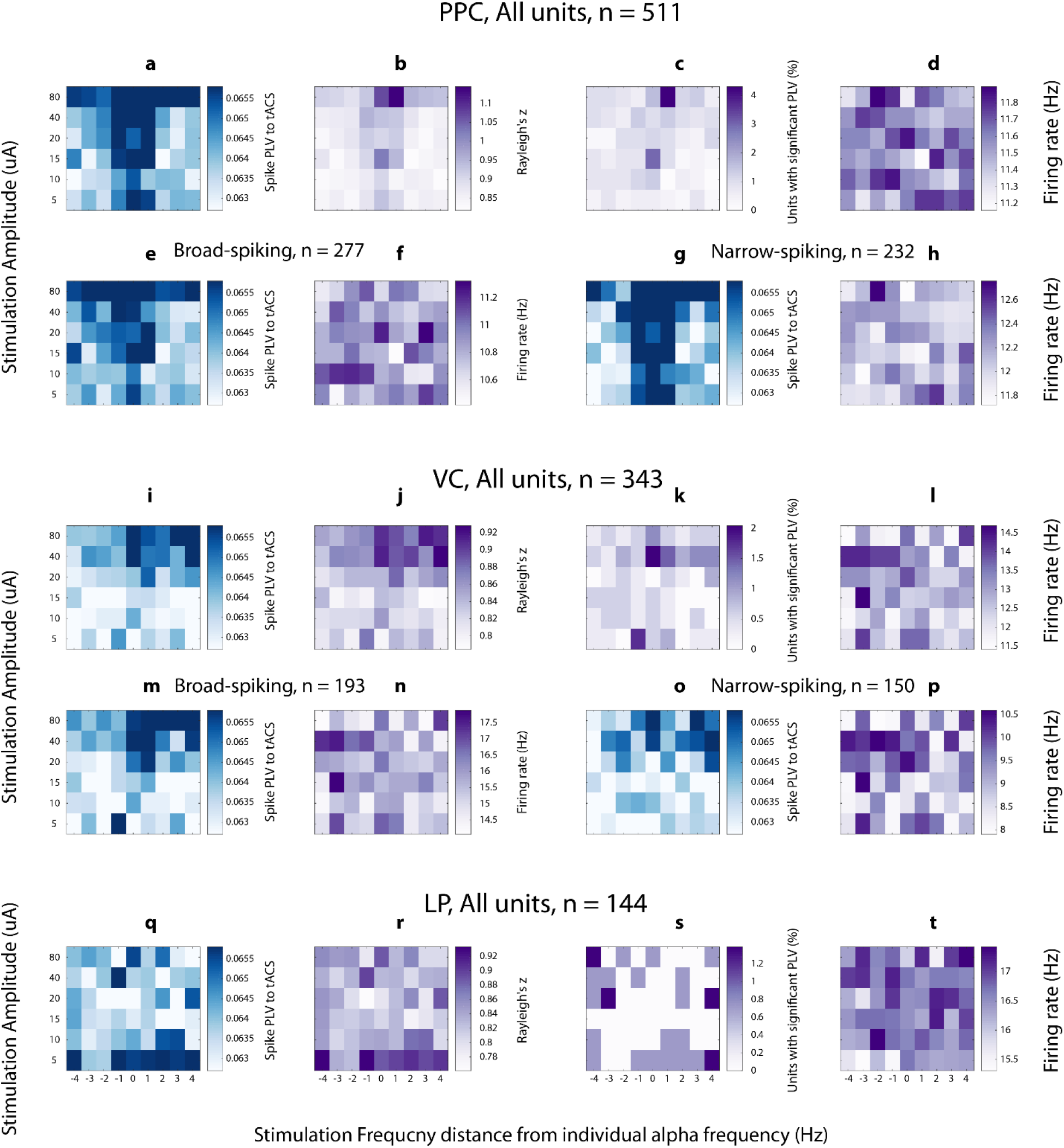
Synchronization and firing rate maps for animal 2. **(a)** Average phase locking of PPC spikes (averaged across 621 units) to tACS phase as measured by phase-locking value (PLV) as a function of stimulation frequency (horizontal axis) and stimulation amplitude (vertical axis). **(b)** Average Rayleigh’s z-score (across all PPC units) as a function of stimulation parameters. **(c)** Percentage of PPC units with significant PLV as indicated by Rayleigh’s test. **(d)** Average firing rate of all PPC units. **(e)** Average PLV of broad-spiking PPC units (n=382). **(f)** Average firing rate of broad-spiking PPC units. **(g)** Average PLV of narrow-spiking PPC units (n=229). **(h)** Average firing rate of narrow-spiking PPC units. **(i-p)** Corresponding measures of **(a-h)** for VC. **(q-t)** Corresponding measures of **(a-d)** for LP. The horizontal axis shows the distance from endogenous alpha frequency in Hz. Related to Figure 8.

**Figure S4.**
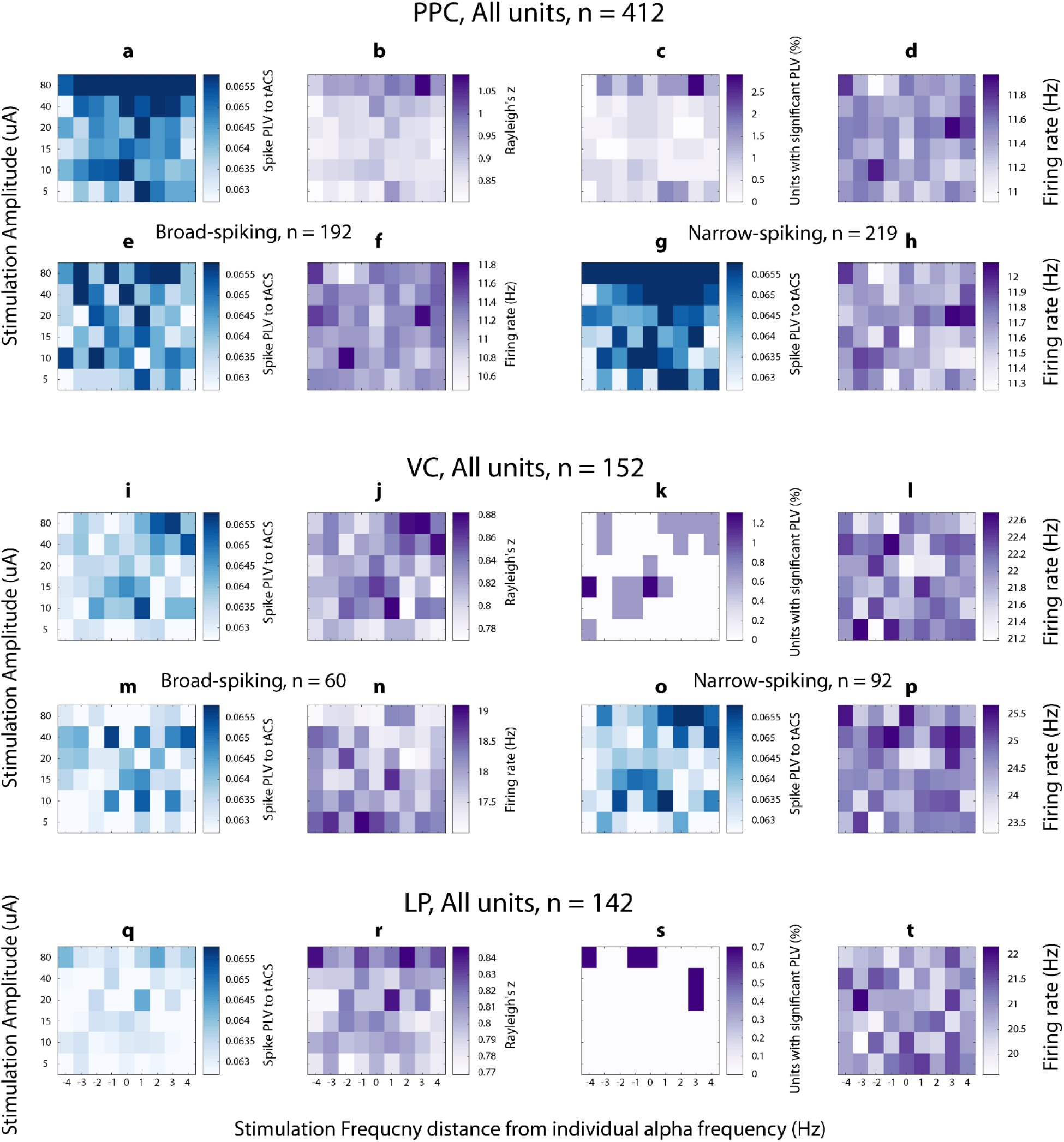
Synchronization and firing rate maps for animal 3. **(a)** Average phase locking of PPC spikes (averaged across 621 units) to tACS phase as measured by phase-locking value (PLV) as a function of stimulation frequency (horizontal axis) and stimulation amplitude (vertical axis). **(b)** Average Rayleigh’s z-score (across all PPC units) as a function of stimulation parameters. **(c)** Percentage of PPC units with significant PLV as indicated by Rayleigh’s test. **(d)** Average firing rate of all PPC units. **(e)** Average PLV of broad-spiking PPC units (n=382). **(f)** Average firing rate of broad-spiking PPC units. **(g)** Average PLV of narrow-spiking PPC units (n=229). **(h)** Average firing rate of narrow-spiking PPC units. **(i-p)** Corresponding measures of (a-h) for VC. **(q-t)** Corresponding measures of **(a-d)** for LP. The horizontal axis shows the distance from endogenous alpha frequency in Hz. Related to Figure 8.

**Figure S5.**
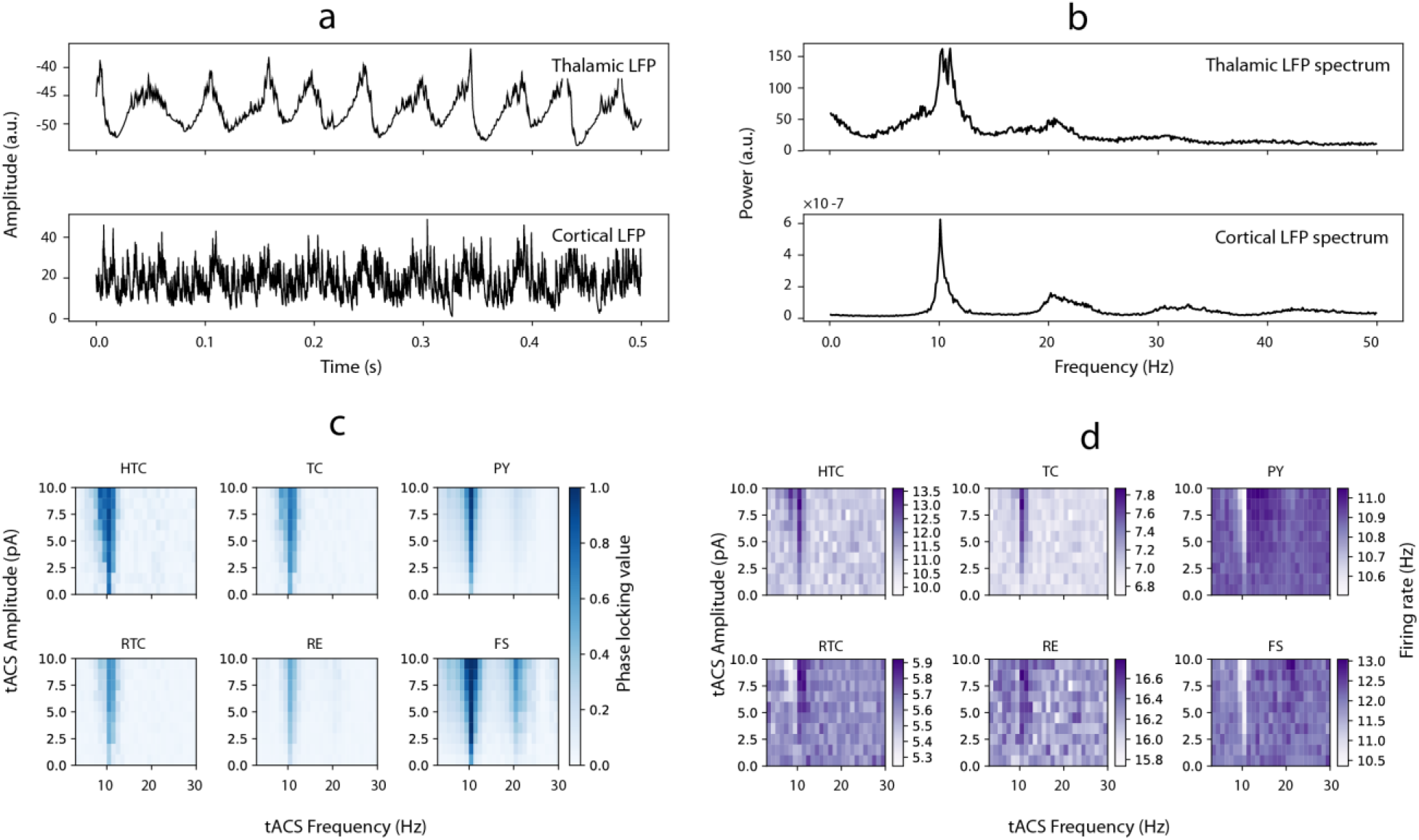
Synchronization of altered thalamo-cortical network by tACS. In the altered network both thalamic and cortical regions oscillates at endogenous alpha instead of theta, the tACS entrains both cortical and thalamic neuronal populations. (a) Both cortical and thalamic LFPs show alpha endogenous oscillations. (b) Color-coded phase locking value (PLV) of spiking activity of different neuron types to tACS as a function of tACS frequency (horizontal axis) and tACS amplitude (vertical axis). Triangular-shape high-synchrony Arnold tongue regions centered on alpha frequency are evident for all neuron types. (c) Color-coded firing rate maps corresponding to PLV maps for all neuron types. Related to Figure 9.

**Figure S6.**
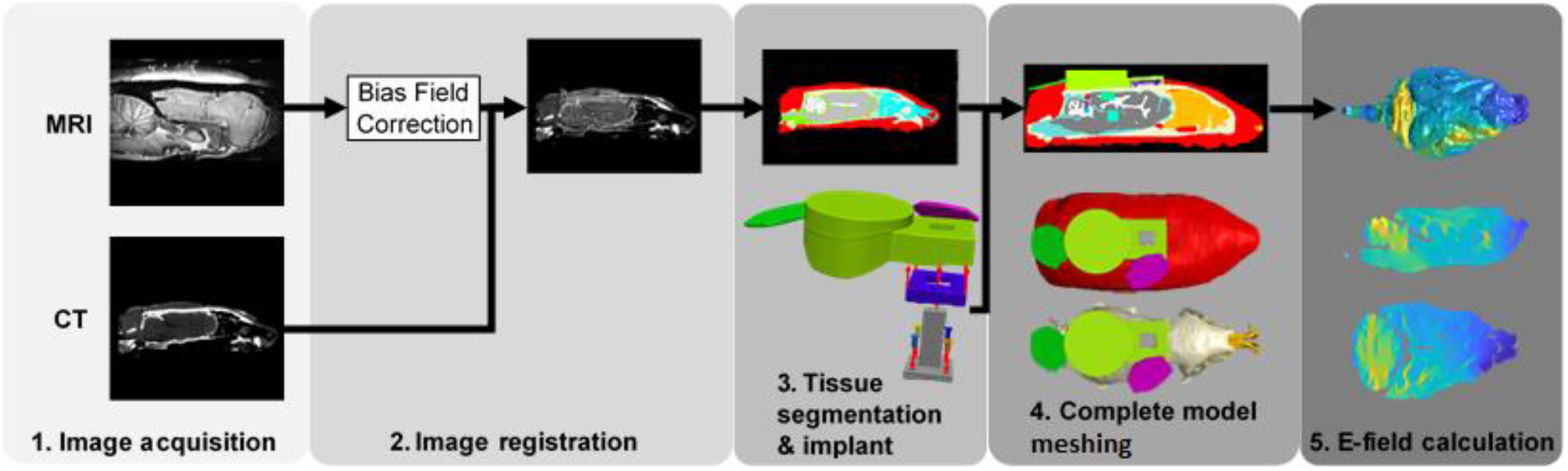
Pipeline for computational modeling for tACS electric field in ferret head. The CT and MRI imaging data were co-registered to allow segmentation of hard and soft tissues (1-3). Implanted hardware and surface tACS electrodes were added to this segmentation (3) and meshed (4) with finite elements to create a three-dimensional representation of the experimental setup. Tissue and hardware conductivities were assigned based on the literature (see Table S1) and COMSOL was used to solve the quasi-static Poisson equation for nodal electric potentials, from which the electric field was calculated (5). Related to Figure 7.

**Table S1.** Tissue/Material electrical conductivities. Related to Figure 7.

| *Tissue/Material* | *Electrical*<br>*Conductivity [S/m]* |
| --- | --- |
| Scalp | 0.29 |
| Skull | 0.03 |
| Cerebrospinal fluid | 1.65 |
| Grey matter | 0.82 |
| White matter | 0.38 |
| Stimulation pad | $5.9 \times 10^7$ |
| Head post, bone screws | $1.39 \times 10^6$ |
| Acrylic glue | $3.6 \times 10^{-5}$ |
| Dental cement | $2.5 \times 10^{-14}$ |
| Air | $2.5 \times 10^{-14}$ |

